# Addition of Multiple Introns to a Cas9 Gene Results in Dramatic Improvement in Efficiency for Generation of Gene Knockouts in Plants

**DOI:** 10.1101/2020.04.03.023036

**Authors:** Ramona Grützner, Patrick Martin, Claudia Horn, Samuel Mortensen, Erin J. Cram, Carolyn W. T. Lee-Parsons, Johannes Stuttmann, Sylvestre Marillonnet

## Abstract

The recent discovery of the mode of action of the CRISPR/Cas9 system has provided biologists with a useful tool for generating site-specific mutations in genes of interest. In plants, site-targeted mutations are usually obtained by stably transforming a Cas9 expression construct into the plant genome. The efficiency with which mutations are obtained in genes of interest can vary considerably depending on specific features of the constructs, including the source and nature of the promoters and terminators used for expression of the Cas9 gene and the guide RNA, and the sequence of the Cas9 nuclease itself. To optimize the efficiency with which mutations could be obtained in target genes in *Arabidopsis thaliana* with the Cas9 nuclease, we have investigated several features of its nucleotide and/or amino acid sequence, including the codon usage, the number of nuclear localization signals (NLS) and the presence or absence of introns. We found that the Cas9 gene codon usage had some effect on Cas9 activity and that two NLSs work better than one. However, the most important impact on the efficiency of the constructs was obtained by addition of 13 introns into the Cas9 coding sequence, which dramatically improved editing efficiencies of the constructs; none of the primary transformants obtained with a Cas9 lacking introns displayed a knockout mutant phenotype, whereas between 70% and 100% of primary transformants generated with intronized Cas9 displayed mutant phenotypes. The intronized Cas9 was also found to be effective in other plants such as *Nicotiana benthamiana* and *Catharanthus roseus*.

## Introduction

The discovery of the mode of action of the CRISPR/Cas9 system in 2012 has provided biologists with a useful tool for generating site-specific mutations in genes of interest in living organisms (Cong et al., 2013; Jinek et al., 2012; Mali et al., 2013). The enzymes of several CRISPR systems including Cas9 and Cas12/Cpf1 have since been used for generating site-targeted mutations in a wide range of organisms that includes animals, plants, fungi and bacteria. In plants, site-specific mutations are generally obtained by transforming plants with a construct that contains the Cas9 gene and a guide RNA expression cassette. Cells from the transformed plants expressing all components of the CRISPR/Cas9 system undergo cleavage of chromosomal DNA at the target site(s). The cleaved DNA is then usually repaired by non-homologous end-joining, resulting in mutated target site sequences. The Cas9 construct can be eliminated from the genome of the transformed plants in the next generation by Mendelian segregation. In plants, like in other organisms, not all transformants containing a Cas9 and guide RNA construct display mutations in the target gene, with efficiencies reported to vary from a few percent to close to 100% (Ahmad et al., 2020; Belhaj et al., 2013). The efficiency with which mutations are generated in target genes depends on multiple factors, including the choice of the target sites in selected genes and the nature of the coding and regulatory sequences of the Cas9 gene and of the guide RNA construct. In recent studies, several parameters of the architecture of Cas9 constructs designed for plants have been investigated, including the codon usage of the Cas9 gene, the number of NLS in the Cas9 enzyme, and the nature of promoters and terminators of the Cas9 gene, the length and sequence of the conserved region of the guide RNA and the terminator sequence of the guide RNA, and the relative orientation of the various expression cassettes in the final T-DNA (Castel et al., 2019; Ordon et al., 2019). These studies suggested that several features of the construct, such as the nature of the promoters driving Cas9 expression and the relative orientation of the guide RNAs and Cas9 gene cassettes, had some effect on the efficiency of the constructs. However, the most significant effect on efficiency was attributed to the nature of the coding sequence of the Cas9 gene. Castel and coworkers compared four different Cas9 coding sequences and found that the most efficient one consisted of a plant codon-optimized sequence that contained one intron and two NLSs (Li et al., 2013). However, since the plant codon-optimized sequence was not tested without an intron, and was also tested only with the two NLSs, it was not possible to conclude how much any of these features individually contributed to the high activity of this particular sequence.

In this work, we have investigated the role of the codon usage, the numbers of NLSs, and the presence of introns in the coding sequence of the Cas9 on genome editing efficiencies in stable transgenic plants. Introns have been known for many years to have a positive effect on gene expression (Callis et al., 1987). For example, introduction of multiple introns in a TMV viral vector construct transiently delivered to *Nicotiana benthamiana* leaves as a T-DNA by *Agrobacterium* led to a large increase in expression of a recombinant protein from the viral vector (Marillonnet et al., 2005). The improvement was found to increase with the number of introns introduced, with the highest efficiency obtained when 12 introns were used. This beneficial effect of introns was hypothesized to be due to an increased efficiency of processing and export of the pre-mRNA viral vector transcript from the nucleus to the cytoplasm of the host cells.

Here, we found that presence of two NLSs in Cas9 improves the efficiency with which mutations can be obtained, but the largest effect was obtained by addition of 13 introns to the coding sequence of Cas9. With the intronized Cas9, single and double knockouts or large chromosomal deletions (30 – 70 kb) were obtained with high frequencies (> 70% and ~ 10%, respectively) in primary Arabidopsis transformants. In a companion manuscript (Barthel et al.), we furthermore demonstrate the mutagenic capacity of the optimized Cas9 gene by generating duodecuple (12x) mutant Arabidopsis plants in a single step. The intronized Cas9 was shown to also work well in *Nicotiana benthamiana* and *Catharanthus roseus*.

## Results

### Comparison of Cas9 genes

Seven Cas9 expression constructs were made with different Cas9 coding sequences. The first Cas9 coding sequence (hCas9, Figure 1 and Figure S1) is a human codon-optimized sequence (Mali et al., 2013). This sequence contains a C-terminal nuclear localization sequence (NLS). We used the domesticated version of this gene (lacking internal type IIS restriction site (Castel et al., 2019; Nekrasov et al., 2013) for use with the modular cloning system MoClo (Weber et al., 2011). This Cas9 sequence was subcloned without or with an additional N-terminal NLS to make Cas9 expression constructs pAGM51511 and pAGM51613. To check the effect of codon-optimization, a second Cas9 coding sequence was synthesized using the *Zea mays* codon usage (zCas9), which has a high GC content of 55% (https://www.kazusa.or.jp/codon/cgi-bin/showcodon.cgi?species=4577). A high GC content codon usage was selected as we planned to introduce introns into this sequence, and GC rich exon sequences can induce more efficient splicing of inserted introns (Carle-Urioste et al., 1997). The codon-optimized Cas9 sequence includes a C-terminal nuclear localization sequence (NLS) with exactly the same amino acid sequence as the hCas9 gene. This Cas9 sequence was also subcloned without or with an additional N-terminal NLS to make two Cas9 expression constructs, pAGM51523 and pAGM51535. A third Cas9 sequence version (zCas9i) was made by introducing 13 Arabidopsis introns into the *Zea mays* codon-optimized version. This Cas9 sequence was also subcloned without or with an additional N-terminal NLS to make two Cas9 expression constructs, pAGM51547 and pAGM51559. Finally, a fourth Cas9 version was made, zCas9io, similar to the intron-containing Cas9, but containing mutations at 4 sites in introns 1, 3, 12 and 13 to remove a potential cryptic splice site in the first intron and to improve the splicing efficiency of some weak splice sites, as predicted by the NetGene2 intron splice site prediction software http://www.cbs.dtu.dk/services/NetGene2/) (Figure S2). This Cas9 version also contains an N-terminal FLAG tag followed by a nuclear localization signal. It was subcloned to make a Cas9 expression construct, pAGM51561. The Cas9 coding sequences in all constructs were cloned under control of the Arabidopsis RPS5a promoter, which drives effective levels of expression of the Cas9 nuclease in Arabidopsis (Ordon et al., 2019; Tsutsui and Higashiyama, 2017).

**Figure 1:**
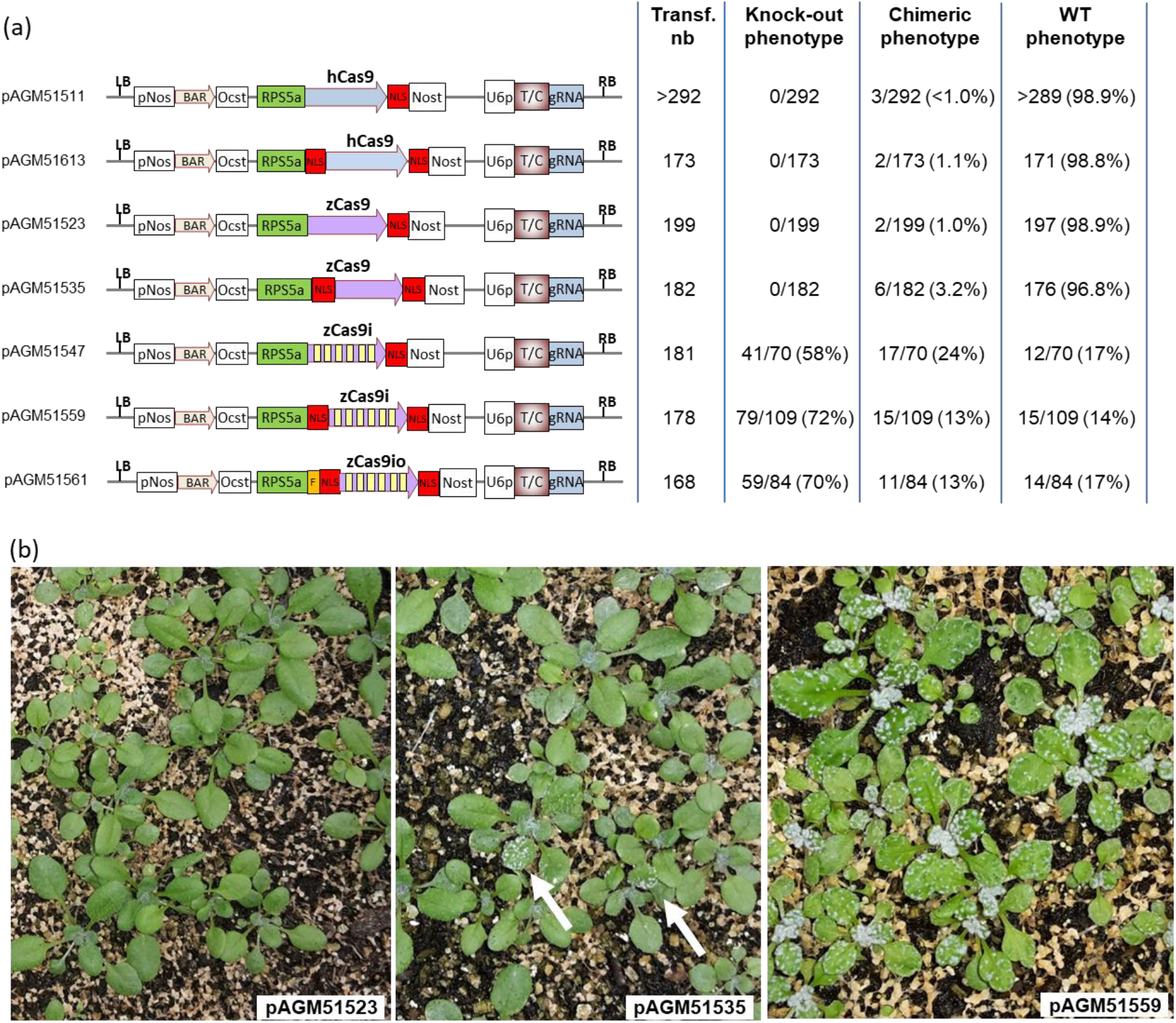
Comparison of different Cas9 versions by mutagenesis of TRY and CPC. **(a)** Structure of the Cas9 constructs and mutagenesis efficiency. The number of transformants obtained for each construct is shown (Transf. nb). For the constructs with intronized Cas9, which produced a lot of plants with mutant phenotypes, transformants from a part of the trays were uprooted to evaluate their phenotypes (70 to 109 randomly selected transformants). pNos: Nopaline synthase promoter; Bar: Bar gene coding sequence; Ocst: Octopine synthase terminator. Rps5a: Arabidopsis ribosomal protein 5a promoter; hCas9: human codon-optimized Cas9 coding sequence; zCas9: *Zea mays* codon-optimized Cas9 coding sequence; zCas9i: the same sequence with 13 introns represented as 6 yellow boxes; zCas9io: sequence variant of zCasi; F: flag tag; Nost: Nopaline synthase terminator; U6p: Arabidopsis U6 promoter; T/C: target sequence of the guide RNA for the *TRY* and *CPC* genes; gRNA: represents the conserved region of the guide RNA. LB and RB: left and right T-DNA borders. **(b)** Picture of the primary transformants of two non-intronized Cas9 constructs pAGM51523, pAGM51535 and one intronized Cas9 construct pAGM51559.

To test the different Cas9 genes, we used a single guide RNA that targets the genes of two homologous Arabidopsis transcription factors, TRY and CPC. These genes act as negative regulators of trichome development and provide a convenient visual marker for Cas9 activity, with knockout plants displaying an increased number of trichomes in leaves (Wang et al., 2015). The guide RNA was cloned under control of a consensus sequence of Arabidopsis U6 promoters (Nekrasov et al., 2013).

The constructs were transformed in Arabidopsis using flower dip transformation, and more than 160 transformants were obtained for each construct. The constructs containing Cas9 without introns did not produce a single primary transformant with a complete knockout phenotype. A few plants displayed some leaf sectors with the expected mutant hairy phenotype (Figure 1; Figure S3). The highest number of plants with such chimeric phenotype was obtained with the *Zea mays* codon-optimized Cas9 with two NLSs (6/182 transformants or 3.2%). The other constructs with Cas9 without introns led to approximately 1% of plants with a chimeric mutant phenotype (7/664), with all other plants displaying a WT phenotype. In contrast, transformation of the intronized Cas9 constructs led to a large number of primary transformants with full knockout phenotypes (Figure 1; Figure S3). Constructs containing the *Zea mays* codon–optimized Cas9 with two NLSs, pAGM51559 and pAGM51561, led to 72% (79/109) and 70% (59/84), respectively, of transformants displaying a full knockout phenotype. The intron-containing Cas9 with a single NLS produced 58% (41/70) of plants with a knockout phenotype.

### Analysis of the transformants

To understand why some plants displayed a mutant phenotype, but not all plants, DNA was extracted from primary transformants of pAGM51559 and pAGM51561 with a wild-type phenotype (8 plants for each construct). In addition, for both constructs, DNA was extracted from 2 plants with a full knockout phenotype and from 4 plants displaying a chimeric phenotype. For these chimeric plants, two DNA extractions were made, one from leaves showing a WT phenotype, and one from leaves with a mutant phenotype. All DNA preparations were analyzed by PCR with two pairs of primers in the Cas9 constructs, one for amplification of the guide RNA region (primers guia1 and 4, product of 583 bp), and one for amplification of a region within the Cas9 sequence (primers casan2 and 3, product of 966 bp). Of the 8 pAGM51559 transformants with WT phenotype, one did not contain the Cas9 and the guide RNA, and probably contained a truncated T-DNA (Figure S4). The other 7 plants with a WT phenotype appeared to contain a full T-DNA. For the 8 pAGM51561 transformants, one contained an incomplete T-DNA containing the Cas9 gene but lacking the guide RNA region, while the other 7 plants appeared to contain a full T-DNA. All other plants, mutant and chimeric, contained both Cas9 sequences and the guide RNA.

All DNAs prepared from the pAGM51559 and pAGM51561 transformants were analyzed by PCR with two pairs of primers designed to amplify fragments of *CPC* and *TRY* that include the target site for Cas9. The PCR products were then sequenced (Figure S5). For pAGM51559 transformants, the plant with WT phenotype that lacked a complete T-DNA did not show any mutation by sequencing, as expected. Interestingly, 2 of the 7 remaining plants with a WT phenotype displayed some mutations at the DNA level, both in the *CPC* and *TRY* genes. These can be seen as double peaks in the sequence traces starting at the Cas9 target site (Figure S5a, b). The remaining 5 plants with a WT phenotype did not show any mutations at the DNA level. Analysis of the plants with chimeric phenotype showed that all DNA samples, both from the leaves with WT and mutant phenotypes, displayed some mutations. Finally, as expected, the two transformants with full knockout phenotype, showed mutations at the DNA level both in the *CPC* and *TRY* genes.

To analyze the mutation spectrum in more detail, the PCR products for the *CPC* gene of 5 transformants for pAGM51559 and pAGM51561 were cloned and sequenced. For pAGM51559, the PCR products selected for sequencing were from one transformant with WT phenotype but containing mutations as previously determined by sequencing (named WT2), from one transformant with knock out phenotype (Mut9), and from three transformants with chimeric phenotype (DNA extracted from leaves with WT phenotypes, CWT11, CWT12 and CWT14, Figure S6). Sequences of the PCR product from the plant with WT phenotype, WT2, showed a mix of mutant sequences and WT sequences. The same was found in sequences from chimeric plants extracted from leaves with WT phenotype. Finally, and as expected, analysis of sequences from plants with full knockout phenotype showed the presence of only mutant sequences. In conclusion, plants displaying WT phenotype in the entire plants, or in some sectors, contain some WT sequences that explain the non-mutant phenotype, but some of these plants also contain mutations.

The same analysis was performed with 5 pAGM51561 transformants (Figure S7 and S8). Out of 7 plants with a wild-type phenotype containing a complete T-DNA, 4 plants displayed mutations at the sequence level. Cloning and sequencing of PCR products as for pAGM51559 transformants led to the same results and conclusions. The overall spectrum of mutations includes insertions of one bp at the cleavage site or deletions of 1, 3 or 7 bp.

In summary, transformants from constructs pAGM51559 and pAGM51561 containing 2 NLSs and 13 introns lead to 70 and 72% of primary transformants displaying a full knock out phenotype, respectively, and 13% of transformants displaying a chimeric phenotype. In addition, one quarter to half of the transformants with a WT phenotype display mutations at the DNA level. This means that 89% of pAGM51559 transformants (79 + 15 + 4 estimated plants out of 109) and 92% of pAGM51661 transformants (59 + 11 + 7 estimated plants out of 84) have an active Cas9 that is capable of generating mutations in the primary transformants.

### The amount of Cas9 protein in the nucleus is correlated with the efficiency of mutagenesis

To understand why constructs with introns work better than constructs without introns, and why two NLSs seem better than one, constructs similar to those described in the first experiment (Figure 1) but with Cas9 cloned under control of the 35S promoter were used for transient expression in *Nicotiana benthamiana* leaves by *Agrobacterium*-mediated delivery, and Cas9 protein accumulation was analyzed on Western blots using a Cas9-specific antibody (Figure 2a). Interestingly, in all cases, the amount of Cas9 protein with 2 NLSs was found to be lower than the amount of Cas9 protein with one NLS expressed from corresponding genes (with the same nucleotide sequence, *e.g.* hCas9(1xNLS) *vs.* hCas9(2xNLS)). When comparing accumulation of Cas9 with a single NLS, similar amounts were detected upon expression from genes with different codon usage, but protein amounts appeared mildly enhanced for the intron-optimized zCas9i gene (Figure 2a). This observation was further reinforced by results obtained for Cas9 with 2 NLSs: protein amounts as evaluated by immunodetection clearly increased upon expression from the intron-optimized genes, and there were no marked differences between zCas9i and zCas9io (Figure 2a).

**Figure 2:**
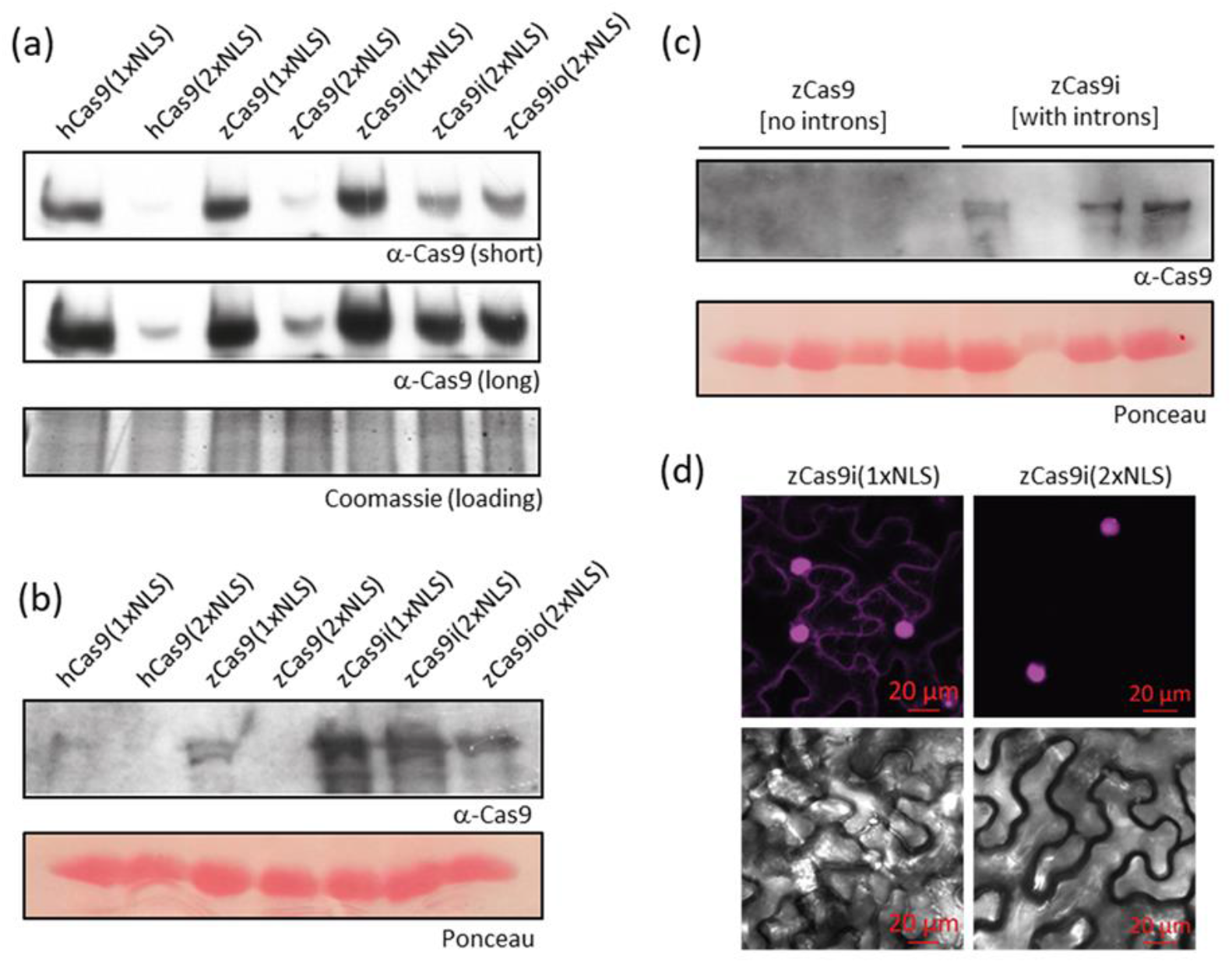
Cas9 nuclease activity correlates with protein accumulation inside the nucleus. **(a)** Accumulation of Cas9 upon expression of different genes in *N. benthamiana*. Indicated Cas9 versions were expressed (under p35S control) in *N. benthamiana* by agroinfiltration. Tissues were used for protein extraction and immunodetection at 3 dpi. The lower part of the PAA gel was stained with Coomassie, and is shown as loading control. **(b)** Accumulation of Cas9 upon expression from different genes in stable Arabidopsis transgenics. Proteins were extracted in pools prepared from leaf tissues of eight independent primary transformants expressing indicated Cas9 versions (with sgRNAs targeting *try* and *cpc*; 5 week-old plants), and used for SDS-PAGE and immunodetection. Pools for the first four samples (Cas9 without introns) were prepared from six wild type-like and two chimeric plants. Pools from transgenics expressing the intron-optimized Cas9 variants were prepared from phenotypically mutant (*try cpc*; hairy) plants. Ponceau staining of the membrane is shown as the loading control. **(c)** Detection of Cas9 in individual T_1_ Arabidopsis transformants. Proteins were extracted from leaf tissues of individual T_1_ transformants expressing either zCas9 or the intron-optimized zCas9i, in both cases with N- and C-terminal NLS signals, and were used for SDS-PAGE and immunodetection. Phenotypically wild type-like tissues were used for zCas9, and *try cpc*-like tissues were used for zCas9i. Ponceau staining of the membrane is shown as loading control. **(d)** Subcellular localization Cas9 versions carrying one or two nuclear localization signals. As in a), but fusions of Cas9 with mEGFP were expressed in *N. benthamiana*, and tissues were used 3 dpi for live cell imaging. Either GFP-Cas9^NLS^ or ^NLS^GFP-Cas9^NLS^ (NLS from SV40) were expressed using the intron-optimized zCas9i gene.

The effect of intron-optimization on protein accumulation was further tested by immunodetection of Cas9 in primary Arabidopsis transformants, either using protein extracts prepared from pooled tissues of 8 primary transformants (Figure 2b), or from individual transformants (Figure 2c). For assembly of pools, 6 plants with WT phenotype and 2 plants with chimeric phenotype were used from transformations with constructs not containing introns, while 8 plants with the mutant phenotype were selected for zCas9i (with 1 or 2 NLSs) and zCas9io. Cas9 levels appeared overall low in Arabidopsis, but immunodetection of Cas9 from pooled samples supported our previous observations from transient expression in *N. benthamiana*: lower protein levels were detected for Cas9 in fusion with 2 NLSs compared to Cas9 with a single NLS, and protein accumulation was enhanced upon expression from the intron-containing genes (Figure 2b). Similarly, we failed to detect Cas9 in protein extracts from individual transformants carrying zCas9(2xNLS) without introns, but the protein was detected in 3/4 plants from transformation of the intron-optimized zCas9i with 2 NLSs (Figure 2c). Taken together, these results suggest that addition of introns to the Cas9 gene leads to a higher expression level of the protein in *N. benthamiana* and Arabidopsis, or potentially also more sustained expression.

Although Cas9 with a single NLS accumulated to higher levels *in planta* in comparison to Cas9 with 2 NLSs, higher mutagenic activities were observed for the latter version (Figure 1). To analyze subcellular localization of Cas9, two constructs coding for GFP-Cas9 fusions (as GFP-zCas9i^NLS^ and ^NLS^GFP-zCas9i^NLS^) under control of the 35S promoter were assembled and used for transient expression in *Nicotiana benthamiana* leaves. The GFP-Cas9 fusion protein containing a single NLS was observed in the nucleus as well as in the cytosol (Figure 2d). In contrast, the version with 2 NLSs was observed predominantly in the nucleus. This suggests that mutagenic activity of Cas9 with a single NLS is limited by inefficient nuclear import. At the same time, presence of 2 NLSs leads to a lower steady-state amount of Cas9 protein, possibly due to enhanced turnover in the nuclear compartment. Intron-optimization of the Cas9 gene appears to counteract this effect by enhancing overall expression, thus ensuring for a nuclease concentration sufficient for efficient mutagenesis directly at the chromatin.

### Test of a construct with low copy number in Agrobacterium

It is useful to be able to recover mutant plants that have lost the Cas9 construct in subsequent generations by segregation. If the T-DNA is present in one copy, one quarter of the mutant plants in the next generation should lack the Cas9 construct. Obtaining plants lacking Cas9 becomes more difficult if several copies of the T-DNA are inserted at different chromosomal locations. It is known that binary vectors that replicate at a single copy in Agrobacterium lead to a higher proportion of transformants containing single copy, backbone-free transgenic plants (Ye et al., 2011). The plasmid backbone used in the experiments described above (Figure 1) has a pVS1 origin of replication, which is known to replicate in Agrobacterium at approximately 20 copies (Heeb et al., 2000; Itoh et al., 1984; Zhi et al., 2015). We therefore tested a binary vector (pAGM37443) that contains the *Agrobacterium rhizogenes* A4 plasmid origin of replication, which replicates in Agrobacterium at approximately 1 copy per cell (Nishiguchi et al., 1987; Ye et al., 2011). The same constructs as described previously (Figure 1) were assembled again in vector pAGM37443 and transformed in Arabidopsis by floral dipping. The number of transformants obtained was smaller than in the previous experiment (12 to 45 transformants for different constructs (Figure 3), compared to > 80 transformants (Figure 1)). Lower transformation efficiencies were caused by seasonal effects and/or plant growth rather than copy number of constructs, as > 170 primary transformants were obtained in an independent transformation repeated later of one of the low copy constructs, pAGM53451 (see below). Nevertheless, the same trend as before was observed: constructs with Cas9 without introns did not produce full knockout phenotypes in the primary transformants, but constructs with intronized Cas9 led to many primary transformants displaying a full knock out phenotype (47.6 and 50%) for constructs with intronized Cas9 and 2 NLSs, Figure 3). The ratio of plants with a full knock out phenotype was slightly lower than when using a pVS1 ori (47% and 50%) for intron-containing constructs and 2 NLS, versus 72% and 70% for the same constructs but with the pVS1 ori. This lower number of knock out plants may be explained by a lower number of plants containing multiple copies of the T-DNA integrated in the genome, as expected from plasmids that replicate at single copy in Agrobacterium. Nevertheless, adding the number of plants with a chimeric phenotype and a complete knockout phenotype, the number of plants carrying mutations at the DNA level is at least 71% (5 + 10 = 15 for a total of 21 plants) and 64% (5 + 18 = 23 for a total of 36 plants) for these two constructs.

**Figure 3:**
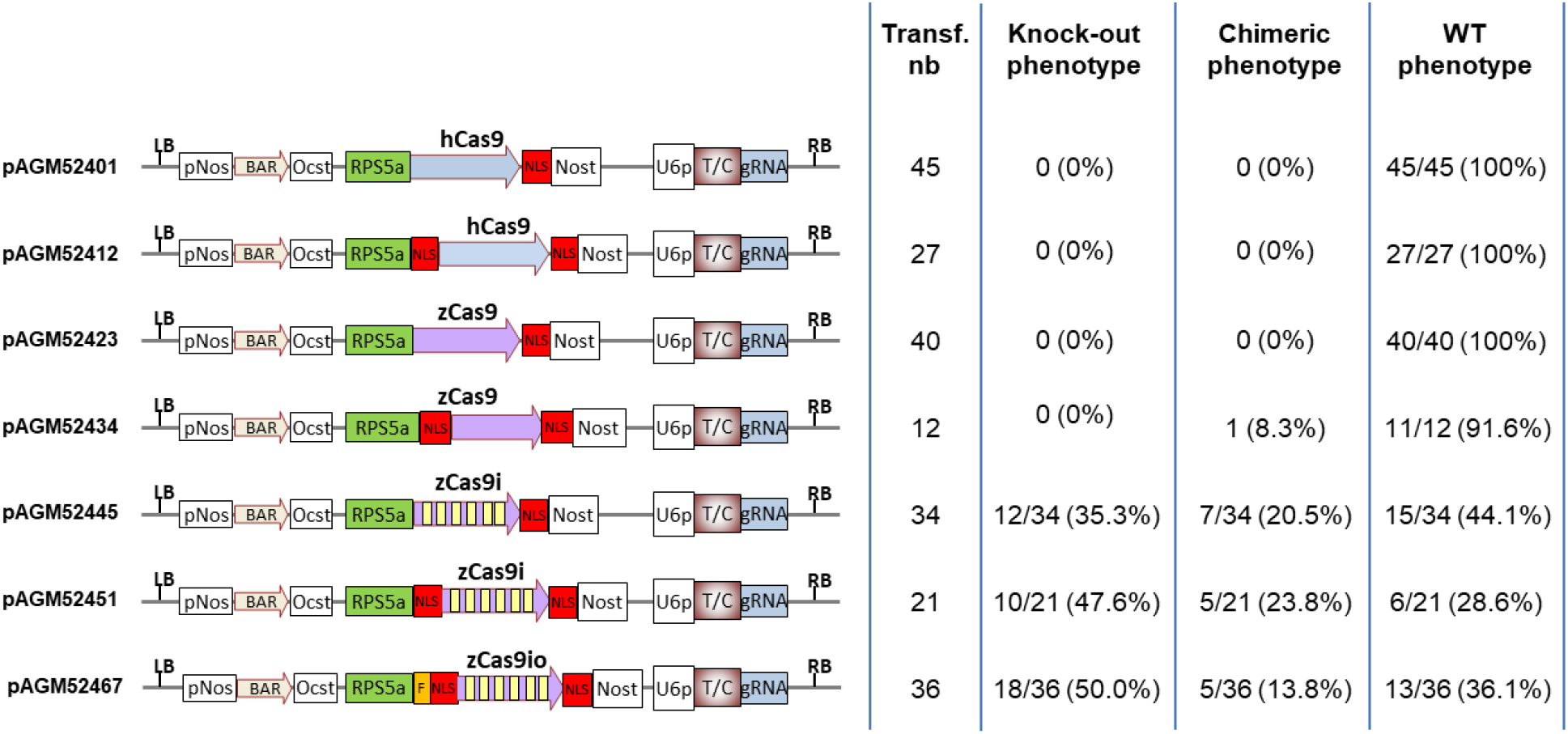
Comparison of different Cas9 versions in low copy vectors by mutagenesis of TRY and CPC. Structure of the Cas9 constructs and mutagenesis efficiency. The legend for the annotations are the same as in figure 1. pNos: Nopaline synthase promoter; Bar: Bar gene coding sequence; Ocst: Octopine synthase terminator. Rps5a: Arabidopsis ribosomal protein 5a promoter; hCas9: human codon-optimized Cas9 coding sequence; zCas9: *Zea mays* codon-optimized Cas9 coding sequence; zCas9i: the same sequence with 13 introns represented as 6 yellow boxes; zCas9io: sequence variant of zCasi; F: flag tag; Nost: Nopaline synthase terminator; U6p: Arabidopsis U6 promoter; T/C: target sequence of the guide RNA for the *TRY* and *CPC* genes; gRNA: represents the conserved region of the guide RNA. LB and RB: left and right T-DNA borders.

Since the first round of transformation did not yield many transformants with the single copy vectors, Arabidopsis transformation was performed again with construct pAGM53451. This time, 174 transformants were obtained, with 53 plants (30.4%) with a wildtype phenotype, 50 plants (28.7%) with a chimeric mutant phenotype and 71 plants (40.8%) with a full mutant phenotype. Therefore, constructs replicating at a single copy in Agrobacterium can produce high number of transformants in Arabidopsis and still produce a high proportion of plants with a chimeric or full knockout phenotype.

### Test of the intronized Cas9 for mutagenesis of genes involved in flower development

To check that significant edits can be obtained in other genes, guide RNAs were made for three genes involved in flower development: *APETALA3* (*AP3;* AT3G54340), *AGAMOUS* (*AG*; AT4G18960) and *LEAFY* (*LFY;* AT5G61850) (Jack et al., 1992; Weigel et al., 1992; Yanofsky et al., 1990). To facilitate assembly of constructs, four cloning vectors were made that contain all components of the final constructs, except the target sequence of the guide RNA (Figures 4a, S9). Two of these vectors contain a BAR cassette for selection of transgenic plants with Basta, and two contain a Kanamycin resistance cassette. The vector backbone contains either the pVS1 ori or the *Agrobacterium rhizogenes* A4 ori. The missing sequence of the guide RNA can be cloned easily by ligating two 24-nucleotide oligonucleotides into the *Bsa*I-digested vector. For the experiment described here, vector pAGM55261 (pVS1 ori) was used (Figures 4a, S9). The constructs for targeting *AP3* (pAGM55361), *LFY* (pAGM55961) and *AG* (pAGM55973) were transformed into Arabidopsis. 86 to 130 transformants were obtained for the three constructs. Before the plants flowered, 12 randomly selected transformants were transferred to single pots. All three transformations gave rise to a large number of plants with a knockout phenotype. All 12 transformants for all three target genes transferred to single pots gave rise to flowering mutant phenotypes. For *AP3*, all 12 plants had a complete knockout phenotype. Counting from the non-transferred plants in the original tray (that were growing in crowded conditions) revealed some plants with WT phenotype (3 out of more than 32 transformants flowering at the time of counting). For *LFY*, 10 plants out of 12 showed a full knockout phenotype and two were chimeric, with a large part of the plants displaying the mutant phenotype, and a few branches only with a WT phenotype that were able to set seeds. For *AG*, 11 plants had a knockout phenotype and one displayed a chimeric phenotype. Summarizing, full mutant phenotypes were observed with a frequency greater than 80% when targeting *AP3*, *AG* or *LFY* with independent constructs each containing one guide RNA and zCas9i. Thus, the intron-optimized Cas9 appears to perform robustly at multiple different loci and with different guide RNAs.

**Figure 4:**
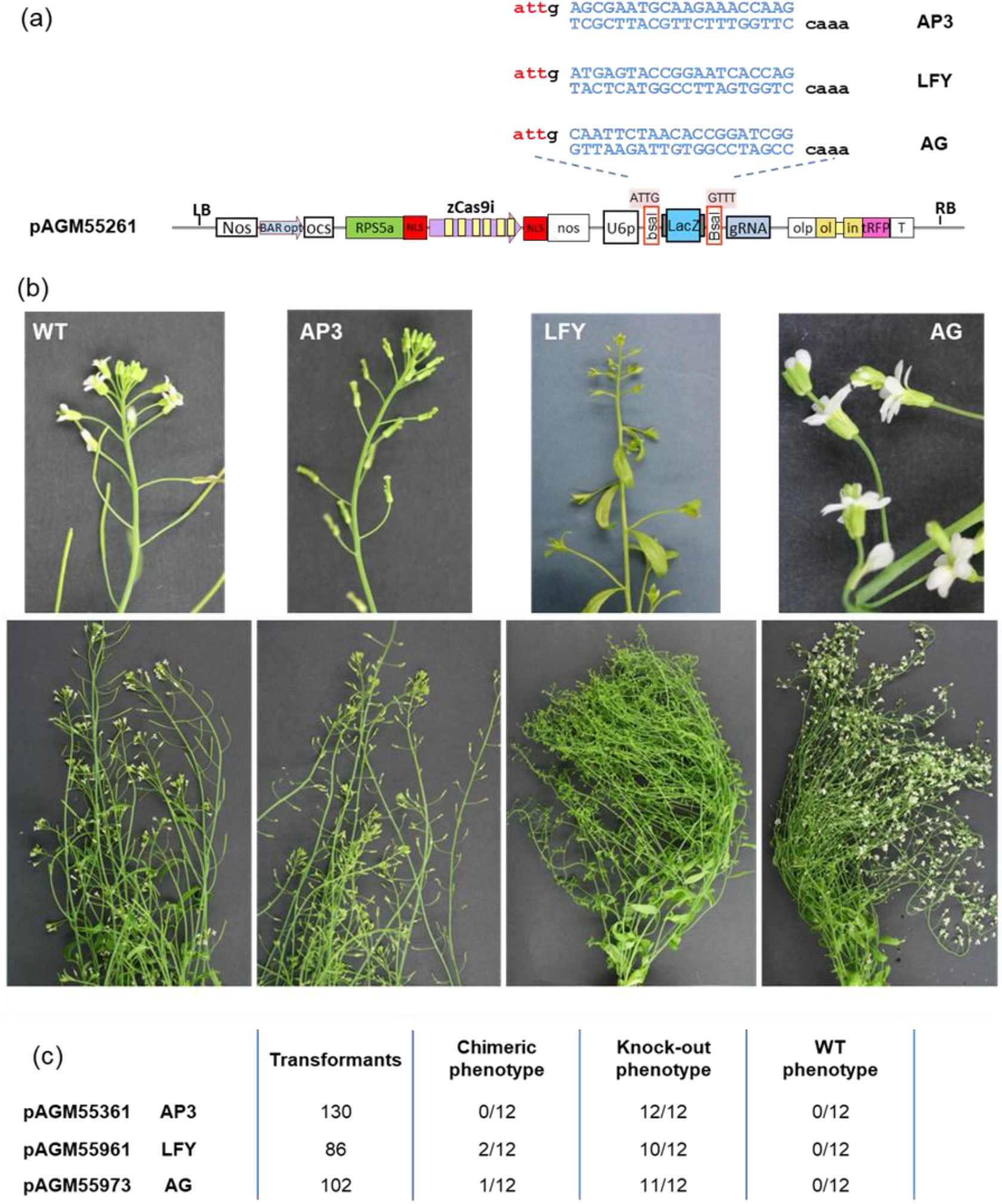
Mutagenesis of floral homeotic genes using intronized Cas9 constructs. **(a)** Schematic representation of the constructs and cloning strategy. For each construct, two oligonucleotides were ligated into the cloning vector pAGM55261 digested with BsaI. The resulting constructs were transformed into Arabidopsis by the floral dip method. **(b)** Phenotypes of the wildtype control and of the transformants. **(c)** Estimation of the number of transformants with wildtype and mutant phenotypes.

### Generation of large chromosomal deletions in Arabidopsis with the intron-optimized Cas9

We previously generated large chromosomal deletions ranging from 70 – 120 kb using hCas9 without introns and under control of a ubiquitin promoter (Ordon et al., 2019; Ordon et al., 2017). In these experiments, deletions occurred at low frequencies (0.5 – 2%), and could be detected in the T_2_ generation only. The efficiency of zCas9i (under *pRPS5a* control) for generation of chromosomal deletions was tested by targeting the *RPP5* and *RPP2* gene clusters (Figure 5). Two different constructs each containing four guide RNAs were assembled (Figure 5a), to program Cas9 for two target sequences on each side of the respective clusters (Figure 5b). Constructs were similar to those used in previous experiments (Figure 1), but also contained the FAST marker (Shimada et al., 2010) for transgene counter-selection by seed fluorescence. Frequencies of deletion events were evaluated directly in primary T_1_ transformants by PCR screening (Figure 5c). Deletions with an expected size of approximately 83 kb were detected with a frequency ~ 10% (4/34) at the *RPP5* locus (Figure 5c). The *RPP2* locus (~ 30 kb) was targeted for deletion both in wild-type Col plants and in the background of a *Δrpp5* mutant line. The expected deletion was detected in 15 – 20% of screened primary transformants (Figure 5e). Furthermore, the inheritability of deletions detected in T_1_ was tested in transgene-free T_2_ segregants, as selected by absence of seed fluorescence (Figure 5d). Chromosomal deletions detected in the T_1_ generation were inheritable for 6/8 families (Figure 5e). Thus, deletions likely occurred early in development, as also observed phenotypically when targeting *try cpc*, and were thus efficiently transmitted to the germline. Taken together, the intron-optimized Cas9 thus induces chromosomal deletions at relatively high frequencies (10 – 20% for deletions ranging up to 80 kb), and deletion lines can easily be isolated from T_2_ segregants of as few as 2 −3 PCR-confirmed T_1_ plants.

**Figure 5:**
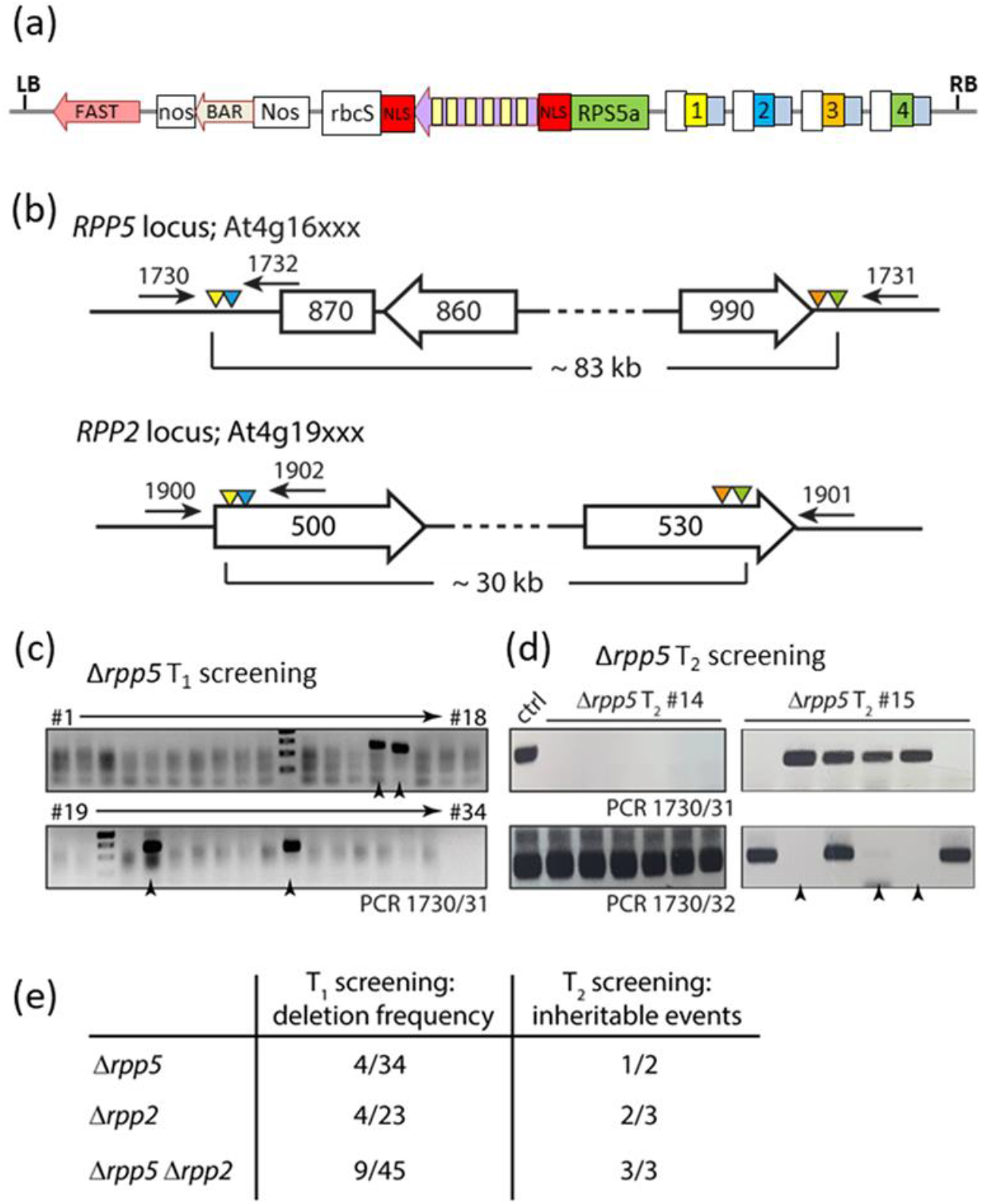
Generation of large chromosmal deletions in Arabidopsis using the intron-optimized zCas9i. **(a)** General architecture of constructs used for generation of large deletions. sgRNAs (under pU6-26 control) are transcribed in the opposite direction relative to zCas9i. FAST - pOle1:Ole1-tagRFP_tole1 (Shimada et al., 2010). **(b)** Schematic drawing of the *RPP5* and *RPP2* loci with sgRNA target sites (triangles; color code corresponding to a) and primer binding sites indicated. Genes are depicted by arrows with Arabidopsis gene identifiers, and a transposable element is depicted by a square. Not drawn to scale. **(c)** Screening of T1 plants for candidate lines carrying a deletion at the *RPP5* locus. T1 plants were selected by resistance to BASTA, and screened by PCR with indicated primers for occurrence of a deletion at the *RPP5* locus. 34 independent T1 lines were screened, putative deletion lines are marked with arrowheads. **(d)** PCR-screening of segregants from T2 families #14 and #15 shown in c) for isolation of a *Δrpp5* deletion line. The upper panel shows a PCR detecting presence of the *Δrpp5* deletion allele (oligonucleotides 1730/31); DNA from a T1 plant was used in the lane marked control (ctrl). The lower panel shows a PCR for detection of the wild type *RPP5* locus (1730/32); Col DNA was used as control. **(e)** Summary on frequencies of large deletions in Arabidopsis when using intron-optimized zCas9.

### Test of the intronized Cas9 in *Nicotiana benthamiana*

The human-codon optimized Cas9 and the intronized zCas9i with 2 NLSs were further tested in *Nicotiana benthamiana.* Four constructs containing either hCas9 or zCas9i together with guide RNAs rt1 or rt4, were designed to target two *Nicotiana benthamiana* genes with putative rhamnosyltransferase activity (Figure S10). With guide RNA rt1, all obtained primary transformants (hCas9: 4/4; zCas9i: 11/11) had mutations at the target site (analyzed by amplification of target site sequences and sequencing of the PCR product). This suggests that both Cas9 genes can mediate genome editing with high efficiencies in *N. benthamiana*, and is in agreement with previous reports on efficient editing in this species with hCas9 (Adachi et al., 2019; Castel et al., 2018; Ordon et al., 2017). In contrast, no mutations were detected in any of the putative transformants (hCas9: 0/7, zCas9i: 0/6) when using rt4 (targeting a different gene), suggesting that the guide RNA for this selected target site did not work.

Eight additional target sites in either rhamnosyl transferase genes or a flavonol synthase gene were tested with the intronized Cas9 only (Figure S10). For one of these (guide fls3), no mutation could be obtained in 14 putative transformants with the intronized Cas9 gene. For the 7 other targets, mutations were obtained in at least some plants, with frequencies ranging from 25 – 100% of primary transformants with mutations (Figure S10). However, these numbers are not fully quantitative, because it was not tested whether putative transformants without mutations in target sites indeed contained a complete T-DNA or were even transgenic.

In addition, induction of small deletions by targeting multiple sites within a target gene was attempted in *N. benthamiana* using zCas9i with two NLSs. In these experiments, a pDGE binary vector with a pVS origin of replication was used (similar to pDGE463; companion manuscript (Barthel et al.)). Three different genes, *Roq1* (*Recognition of XopQ 1*), *NRG1* (*N requirement gene 1*) and an *NPR1* (*Nonexpressor of Pathogensis-Related Genes 1*) homolog were targeted by multiplexing with two (*NRG1*) or three (*Roq1, NPR1*) different sgRNAs (Figure 6a; see experimental procedures for target sites). The distance between the outmost expected cut sites in the targeted genes varied from 57 nt (*NRG1*) to 265 nt (*Roq1*). DNA was extracted from primary transformants, and used for PCR screening using primers flanking target sites (Figure 6b). All three groups of transformants contained several plants with PCR-detectable deletions between target sites (*Roq1* 7/10, *NRG1 7/10*, NPR1 *6/11*). Notably, a PCR product corresponding to size of the wild-type was no longer detectable in several plants due to editing of *Roq1* and the *NPR1* homolog (Figure 6b; red arrows). More than two bands were amplified from several transformants (Figure 6b; blue arrows), suggesting somatic chimerism in the tissues used for DNA extraction. Considering together the data from generation of point mutations and deletions, we conclude that the intronized Cas9 mediates efficient induction of double strand breaks at target sites in *N. benthamiana*.

**Figure 6:**
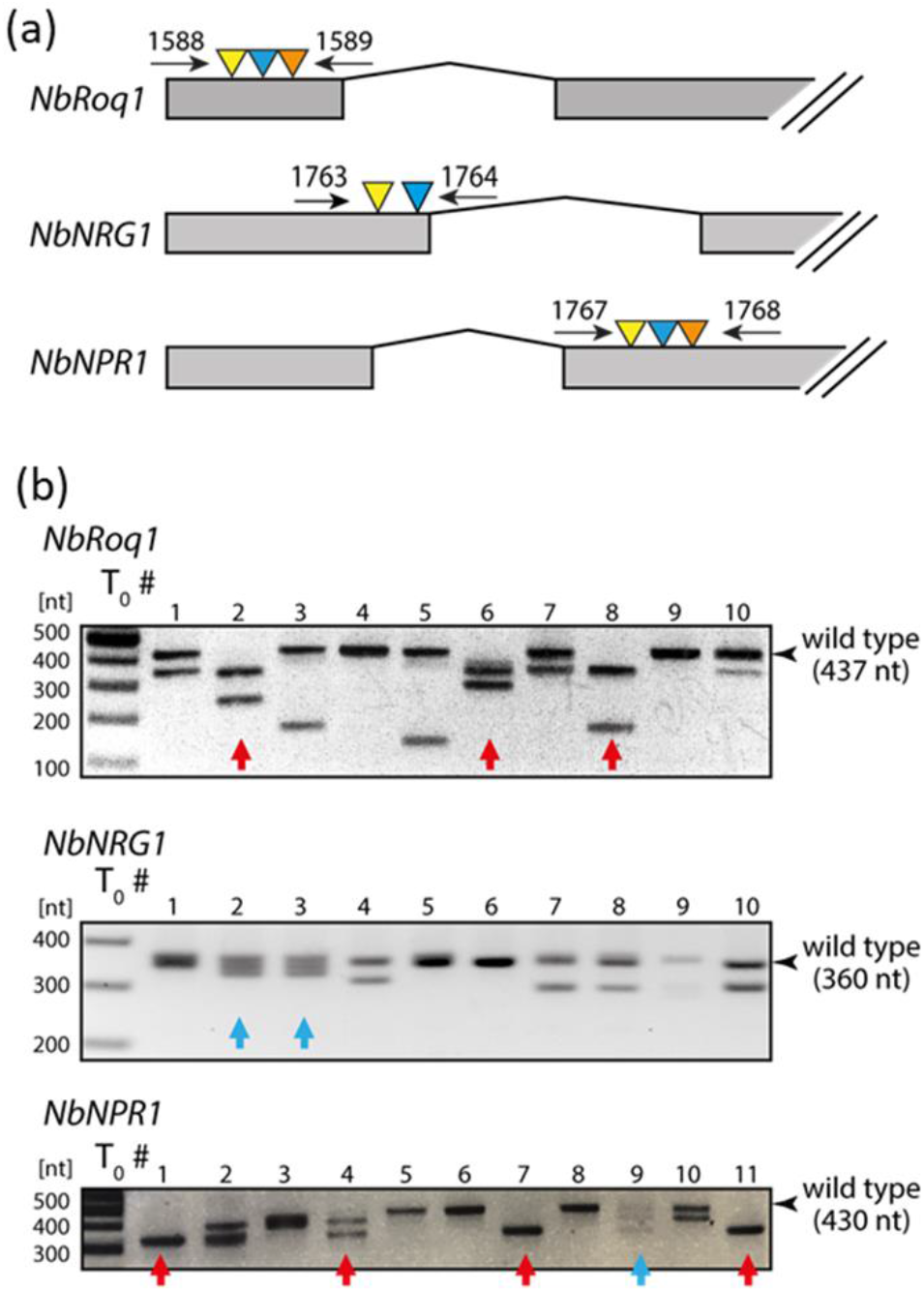
Frequency of small deletions in *N. benthamiana* using zCas9i. **(a)** Gene models of loci targeted for induction of deletions by multiplex genome editing in *N. benthamiana*. Constructs used for editing depicted loci were of similar architecture as shown in Figure 5a, but zCas9i expression was driven by the 35S promoter. *Roq1*: GenBank entry MF773579.1 (Schultink et al., 2017); *NRG1*: Niben101Scf02118g00018, GenBank entry DQ054580.1; *NPR1*: NPR1-like gene, Niben101Scf14780g01001. **(b)** PCR-genotyping of primary (T_0_) transformants. For each transformation, results from 10-11 randomly chosen T_0_ individuals are shown, and genotyping was conducted using oligonucleotides indicated in a). Red arrows mark individuals lacking a PCR product corresponding to the wild type fragment. Blue arrows mark individuals with multiple (> 2) PCR products, indicative of somatic chimerism.

### Gene knockout in hairy roots of *Catharanthus roseus*

The intronized Cas9 was also tested in another plant species, the Madagascar periwinkle, *Catharanthus roseus,* to mutate and delete genes three genes, the Jasmonate-Associated MYC2 (JAM) transcription factors, *JAM2* and *JAM3*, as well as Repressor of MYC2 Targets 1, *RMT1* (Figure 7 and S11). Two constructs were made using the intronized Cas9 with 2 NLS (zCasi) cloned under control of an Arabidopsis ubiquitin promoter (*AtUBQ10*) in a pDGE vector (pDGE454; pVS1 ori). The first construct, pSB311, contains 8 guide RNAs (under pU6-26 promoter control) targeting 2 genes (*JAM2* and *JAM3*), while the second construct, pSB312, contains 8 guide RNAs targeting 3 genes (*JAM2*, *JAM3*, and *RMT1*). The third construct, pSB310, served as control, expressing the intronized Cas9 (zCasi) but not containing sgRNAs. After *Agrobacterium-*mediated transformation and selection with hygromycin B, independent transgenic hairy roots lines were generated (30 lines with pSB311, 19 lines with pSB312, 26 lines with pSB310), and the best growing lines moved from plates into liquid culture. A subset of these hairy root lines was tested for transgene integration: 19/19 were positive for the hygromycin B resistance gene and 14/19 had the guide RNA region.

**Figure 7:**
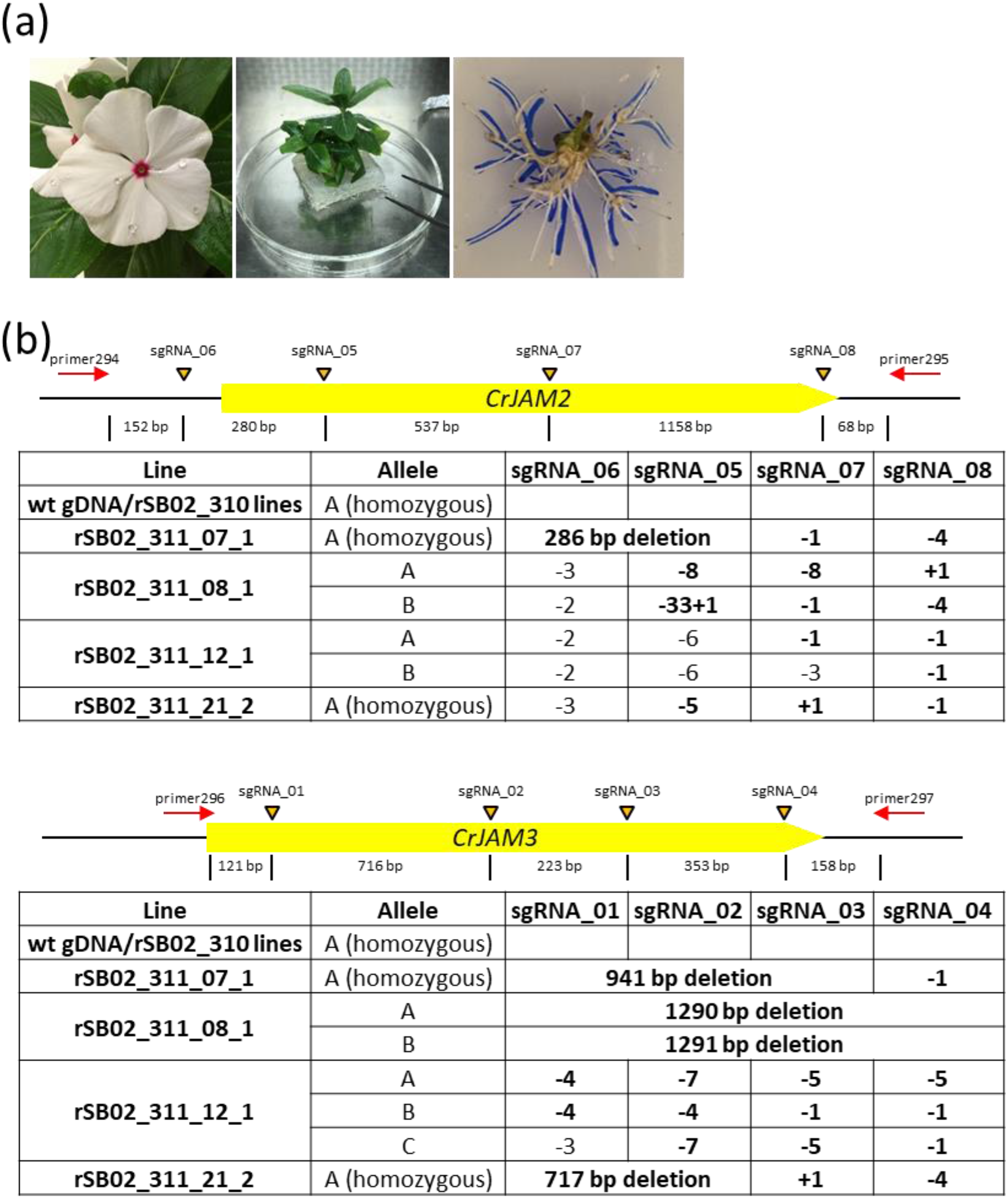
Site-targeted mutagenesis in Catharanthus roseus using the intronized Cas9. **(a)** Picture of wildtype plants and transgenic roots. **(b)** Summary of mutations generated by multiplexing in transgenic hairy root lines in *CrJAM2* and *CrJAM3*.

From the confirmed lines, four lines for each construct (pSB311 and pSB312) were further characterized for mutations in the genes of interest. If different sized PCR products were obtained for a line, these PCR products were directly sequenced. If only one PCR product was obtained, it was sequenced to see if it contained different mutations, and if different mutations were observed, the different products were cloned and individual colonies were sequenced to identify the different mutations. In total, 8 independent lines were tested for 2 or 3 alleles at eight target sites. We observed 100% editing at the target sites (Figures 7, S11). Some lines seemed to be homozygous, and one line appeared chimeric with three different alleles. The spectrum of mutations includes insertion of 1 nucleotide, deletions from 1 to 64 nucleotides, and deletion of all sequences between some pairs of sgRNAs. Two control lines, containing the Cas9 construct but lacking guides, were also sequenced for the gene of interest region and no mutations were observed. Thus, zCas9i was also highly efficient for multiplex editing in *C. roseus* hairy roots, with an apparent mutagenesis efficiency of 100%.

## Discussion

We show here that the codon composition of the Cas9 sequence, the number of encoded NLSs and the presence of introns in the coding sequence can have an impact on the expression level of the Cas9 nuclease and on the final DNA cleavage efficiency in transformants. In our study, the effect of codon-optimization on Cas9 nuclease activity was, however, only minor. Two Cas9 genes with either human or maize codon-optimized sequences were compared (Figure 1a). Both versions (lacking introns) displayed very low activity, with no full knockout phenotypes observed in primary transformants and only a few of the transformants showing mutant leaf sectors. With the Cas9 versions with two NLSs, such chimeric plants were obtained in slightly higher frequencies with the maize codon-optimized version than with the human-codon optimized version (Figure 1). A small effect of codon-optimization on Cas9 activity was also reported in a study that compared the same human codon-optimized version that we used in this work with an Arabidopsis codon-optimized Cas9 (Castel et al., 2019). It is, however, likely that codon-optimization alone may in some cases be sufficient to increase the efficiency in target organisms, as codon-optimized Cas9 genes have been used successfully in several species, such as rice (Mikami et al., 2015; Zhou et al., 2014).

Nuclear import of the Cas9 nuclease is critical for cleavage of chromosomal DNA in eukaryotic cells. In animal systems, a substantial improvement in Cas9 nuclear localization and also activity has been reported for Cas9 versions decorated with up to four NLSs (Koblan et al., 2018; Maggio et al., 2020). In plants, many versions of Cas9 with either one or two NLSs have been reported to work in genome editing (Belhaj et al., 2013), but importance of nuclear import was not analyzed in detail. In this work, constructs with the same Cas9 sequence containing either one or two NLSs were systematically compared to evaluate the impact of the presence of one or two NLSs. When using Cas9 genes without introns, the efficiency for obtaining plants with a chimeric phenotype increased slightly from 1% to 3.2% with one and two NLSs, respectively (Figure 1a). Also, a construct with the intron-containing maize codon-optimized sequence with 2 NLSs (pAGM51559) led to 72% of plants with a knock-out phenotype in primary transformants in comparison to only 58% with a similar construct with only one NLS (pAGM51547, Figure 1). The same trend was also observed with similar constructs in low copy vectors (pAGM52541 and pAGM52445, Figure 3). Interestingly, transient expression of the Cas9 gene in *N. benthamiana* and analysis of Cas9 accumulation in primary Arabidopsis transformants indicated that the steady state amount of Cas9 in cells was lower for a version that has two NLSs (Figure 2a,b). This suggests that the turnover rate of Cas9 in the nucleus may be higher than in the cytoplasm. It is known that the turnover of some proteins that contain degradation signals can be a higher in the nucleus than in the cytosol (Lenk and Sommer, 2000). In yeast, this was linked to higher ubiquitination rates upon nuclear import of the protein. Although Cas9 is not naturally present in eukaryotes, a similar phenomenon may be taking place when the Cas9 protein is transported to the nucleus. We also demonstrated that a GFP-Cas9 fusion containing two NLSs was predominantly observed in the nucleus, while it was observed both in the nucleus and the cytosol with a single NLS (Figure 2d). These results suggest that a single NLS at the C-terminal end is not sufficient for efficient transfer of the Cas9 enzyme to the nucleus, thus limiting nuclease activity at the chromatin.

The most important feature of the Cas9 gene for efficient generation of site-targeted mutations, at least in Arabidopsis, was the presence of multiple introns in the Cas9 coding sequences. This factor alone was sufficient to convert a weakly active Cas9 construct that produced no plants with a knockout phenotype in primary transformants into a very efficient construct that led to between 70% and 100% of transformants displaying a mutant phenotype when targeting one or two genes with a single guide RNA for each (Figures 1, 3, 5; the trichome phenotype used for comparison of the different Cas9 versions depends on inactivation of all copies of the *TRY* and *CPC* genes – four alleles). In a companion manuscript, we furthermore demonstrate the efficiency of the intron-optimized Cas9 by generating duodecuple (12x) Arabidopsis mutants by multiplexing with 24 guide RNAs (Barthel et al.). There are two possible explanations for how intron-optimization of the Cas9 gene might improve editing efficiencies in Arabidopsis. First, intron-optimization leads to higher accumulation of the Cas9 nuclease, as we showed both for transient expression in *N. benthamiana* and stable Arabidopsis transgenics (Figure 2). This effect may be caused by enhanced gene expression and/or nuclear export of the transcript (Shaul, 2017). Additionally, the inclusion of introns into the Cas9 gene may counteract transgene silencing and lead to more sustained expression of the nuclease during development (Christie et al., 2011). Indeed, we rarely observed chimeric plants in previous experiments using mainly hCas9 (Ordon et al., 2019; Ordon et al., 2017), but they regularly occurred with zCas9i. However, it should be noticed that most phenotypes (*e.g.* the hairy phenotype of *try cpc* plants, Figure 1b) occurred early in development, and were already visible in first emerging organs.

While the addition of introns to the Cas9 gene transformed weakly active constructs to efficient ones, it is likely that efficient constructs can be made with Cas9 lacking introns by optimization of the Cas9 sequence alone. Indeed, some Cas9 genes that do not contain introns have been reported to generate high number of transgenic plants with mutant phenotypes (Belhaj et al., 2013; Tsutsui and Higashiyama, 2017). A test of different versions of the nucleases within otherwise identical construct would however be required to make a precise comparison of these sequences.

The presence of introns in the Cas9 sequence may also not be critical for efficient editing in all plant species. For example, the human codon-optimized version that we have used here was reported to work well in some plants such as tomato (e.g. Brooks et al., 2014). Nonetheless, enhanced Cas9 protein accumulation in *N. benthamiana* when using the zCas9i gene suggests that positive effects from intron-optimization are not limited to Arabidopsis, and high mutation rates were obtained in stable *N. benthamiana* lines (Figure 6) and *C. roseus* hairy roots (Figure 7). In conclusion, making an efficient Cas9 construct can be done by inserting multiple introns in the coding sequence, by optimizing the Cas9 gene sequence to lead to high level of Cas9 expression in a given species, or both. Introns may not be necessary for achieving a high expression level but can nevertheless provide a spectacular improvement for the efficiency of the constructs for Cas9 sequences that do not work well enough on their own.

The intron-containing zCas9i performed extremely well in combination with the *RPS5a* promoter (*e.g.* Figure 1a). Surprisingly, we did not obtain genome-edited plants when using zCas9i in combination with the egg cell-specific *EC1.2/DD45* or the ECenh promoter fragments (Castel et al., 2019; Wang et al., 2015) in two independent experiments (data not shown). The intronized Cas9 also worked well with another promoter, the Arabidopsis ubiquitin promoter, in *Catharanthus roseus.* To enable users to make constructs with the intronized Cas9 coding sequence with either the *RPS5a* promoter or other promoters, a series of modules and Cas9 vectors were deposited at Addgene (Figure S9 and S13-16).

In summary, adding introns in a Cas9 gene can significantly improve the efficiency of Cas9 constructs for mutagenesis. Cas9 is also used for other applications such as transcriptional activation or repression. One can speculate that adding introns to these Cas9 versions may have as strong an effect as for use as a nuclease, and potentially even more, as a sustained high level of expression is required for transcriptional repression.

## Experimental procedures

### Plasmid construction

The Cas9 gene sequence was optimized for the codon usage of *Zea mays* using the software from MWG Eurofins. The maize codon usage was used as it is more GC-rich that the codon usage from *Arabidopsis*. The higher GC content is thought to improve intron recognition from exon sequences. Exon/intron splice sites were predicted from the optimized sequences using the NetGene2 online software (http://www.cbs.dtu.dk/services/NetGene2/) (Hebsgaard et al., 1996). For introduction of introns in the Cas9 gene sequence, several Arabidopsis introns were tested *in silico*, and only those that were well recognized by the software after insertion in Cas9 sequences were selected. The Cas9 coding sequence was then constructed using a combination of gene synthesis with DNA gene fragments ordered from MWG Eurofins and of PCR amplification from *Arabidopsis* genomic DNA for the introns. Since the Cas9 sequence is quite large, 6 sub-parts with or without introns were cloned as level −1 modules using Golden Gate cloning and the MoClo system. The subparts were then assembled as level 0 modules containing the entire coding sequence with either one or two NLSs, resulting in modules pAGM12591 and pAGM47539 (*Zea mays* codon-optimized Cas9 without introns and with one or two NLSs, respectively), pAGM13741, pAGM47523 and pAGM51073 (*Zea mays* codon-optimized Cas9 with 13 introns and with one, two and two NLSs, respectively, Figure S1). pAGM47523 and pAGM51073 differ by a N-terminal flag tag in pAGM51073 and by mutations at 4 sites in introns 1, 3, 12 and 13 to try to remove potential cryptic splice sites in nearby exon sequences (Figure S1). The complete sequences of the different Cas9 gene sequences are provided in the supplementary information (Figure S12). The final Cas9 expression constructs in Figure 1 and 3 were made using the MoClo system (Engler et al., 2014; Marillonnet and Grutzner, 2020).

Plasmids containing a Cas9 expression cassette and a transformation marker as well as two BsaI sites for cloning of a single guide RNA (pAGM55261, pAGM55273, pAGM55285 and pAGM55297, Figure S9) are available from Addgene. The intronized Cas9 coding sequence (pAGM47523, zCas9i) is also available from Addgene as a level 0 module. This will provide users with the ability to make constructs with other promoters, and to make constructs with multiple guide RNAs, using the modular cloning system MoClo (https://www.addgene.org/kits/marillonnet-moclo/). The strategy for assembling Cas9 constructs using MoClo is provided in Figures S13 to S16.

### Plant transformation, growth and selection

Constructs were transformed in *Agrobacterium* strain GV3101 pMP90 by electroporation. *Arabidopsis* Columbia-0 (Col) plants were transformed using *Agrobacterium* strains by the flower dip method (Clough and Bent, 1998). Primary transformants were selected by spraying with a BASTA solution (Glufosinate ammonium 0.2 g/l). *N. benthamiana* plants were transformed as previously described (Gantner et al., 2019); a detailed protocol is provided online (dx.doi.org/10.17504/protocols.io.sbaeaie). *C. roseus* hairy roots were generated as described in Rizvi et al. (Rizvi et al., 2015) with the modification made in Mortensen et al., 2019 (Mortensen et al., 2019).

### Guide RNA design

For mutagenesis of *CPC* and *TRY*, a validated guide RNA target sequence was used (Wang et al., 2015): G aatatctctctatctcctc <u>tgg</u> (protospacer adjacent motif, PAM underlined). The first G is not in the target site and serves as the transcription start site of the *Arabidopsis* U6 promoter. Guide RNAs for genes regulating flower morphology (*AP3*, *LFY*, *AG*) and for generation of large deletions (*RPP5* and *RPP2* clusters) were selected using CHOPCHOP (https://chopchop.cbu.uib.no/) (Labun et al., 2019). sgRNA target sites for editing in *N. benthamiana* were selected using CRISPR-P 2.0 (Liu et al., 2017). *C. roseus* sgRNA sequences were selected using Benchling (benchling.com) and on-target scores were predicted (Doench et al., 2016).

### Immunodetection and live-cell imaging

Protein extracts were prepared by grinding tissues in liquid nitrogen and boiling in Laemmli buffer. Total extracts were resolved on 6% SDS-PAGE gels and proteins transferred to nitrocellulose membranes. A rabbit monoclonal α-Cas9 antibody (Abcam EPR18991) was used together with HRP-conjugated secondary antibodies (GE Healthcare), and SuperSignal West Pico and West Femto chemiluminescence substrates (Thermo Fisher) were used for detection. Subcellular localization of GFP-Cas9 variants was determined under a Zeiss LSM780 confocal laser scanning microscope. GFP was excited using the 488 nm laser, and the detector range was set to 493-532 nm.

## Acknowledgements

We thank Nicola Patron and Jonathan D. G. Jones from the Sainsbury lab at the John Innes Centre in Norwich for providing the hCas9 gene module and a level 0 construct containing a consensus sequence of the Arabidopsis U6 promoter (pICSL90001). JS acknowledges Carola Kretschmer and Bianca Rosinsky for excellent technical assistance and greenhouse work. The work of SMa, RG and CH was supported by internal funding of the IPB. Work on Catharanthus roseus was supported by the National Science Foundation (NSF) MCB Award #1516371 to CL-P and EC.

## Supporting Information

Figure S1: Cas9 coding sequence level 0 modules.

Figure S2: Optimization of Zcas9i to improve the predicted splicing efficiency.

Figure S3: Pictures of primary transformants transformed with constructs targeting CPC and TRY

Figure S4: Analysis of Arabidopsis transformants for presence of a T-DNA.

Figure S5: Sequence analysis of TRY and CPC genes in the pAGM51559 transformants

Figure S6: Sequences analysis of cloned PCR products from pAGM51559 transformants

Figure S7: Sequence analysis of CPC in the pAGM51561 transformants.

Figure S8: Sequences analysis of cloned PCR products from pAGM51561 transformants

Figure S9: Vectors for cloning Cas9 constructs.

Figure S10: Cas9 mutagenesis in Nicotiana benthamiana, overview of results.

Figure S11: Analysis of mutations in Catharanthus roseus obtained with construct pSB312.

Figure S12: Nucleotide sequences of the Cas9 and NLS coding sequences

Figure S13: Cloning of a Guide RNA in a MoClo level 1 construct

Figure S14: Construction of Cas9 constructs containing 1 to 4 guide RNAs

Figure S15: Construction of Cas9 constructs containing 5 to 10 guide RNAs.

Figure S16: Cloning 1 to 6 guide RNAs in level 2 vectors already containing Cas9.

## Author contributions

SMa designed the intronized Cas9 gene and the constructs and experiments for Cas9 comparisons and testing in Arabidopsis and Nicotiana benthamiana. JS designed and tested the Cas9-GFP fusion, supervised the Cas9 quantification experiments and designed constructs and experiments for the generation of deletions in Arabidopsis and benthamiana. RG, CH and PM performed experiments. SMo, EJC and CLP designed and performed the *Catharanthus* experiments. SMa wrote the manuscript with contributions from JS and all authors.

