## Supplementary figures for "Addition of Multiple Introns to a Cas9 Gene Results in Dramatic Improvement in Efficiency for Generation of Gene Knockouts in Plants"

Figure S1: Cas9 coding sequence level 0 modules.

Figure S2: Optimization of Zcas9i to improve the predicted splicing efficiency.

Figure S3: Pictures of primary transformants transformed with constructs targeting CPC and TRY

Figure S4: Analysis of Arabidopsis transformants for presence of a T-DNA.

Figure S5: Sequence analysis of TRY and CPC genes in the pAGM51559 transformants

Figure S6: Sequences analysis of cloned PCR products from pAGM51559 transformants

Figure S7: Sequence analysis of CPC in the pAGM51561 transformants.

Figure S8: Sequences analysis of cloned PCR products from pAGM51561 transformants

Figure S9: Vectors for cloning Cas9 constructs.

Figure S10: Cas9 mutagenesis in Nicotiana benthamiana, overview of results.

Figure S11: Analysis of mutations in *Catharanthus roseus* obtained with construct pSB312.

Figure S12: Nucleotide sequences of the Cas9 and NLS coding sequences

Figure S13: Cloning of a Guide RNA in a MoClo level 1 construct

Figure S14: Construction of Cas9 constructs containing 1 to 4 guide RNAs

Figure S15: Construction of Cas9 constructs containing 5 to 10 guide RNAs.

Figure S16: Cloning 1 to 6 guide RNAs in level 2 vectors already containing Cas9.

**Figure S1: Cas9 coding sequence level 0 modules.**

Human codon-optimized sequences are shown in blue. *Zea mays* codon-optimized sequences are shown in orange, and introns are shown in yellow. Nuclear localization signal (NLS) are shown in red. A flag tag (F) is shown in yellow. All modules are flanked by two BsaI sites (sequences ggtctcaAATG and GCTTtgagacc) providing compatibility for DNA assembly with the MoClo system.

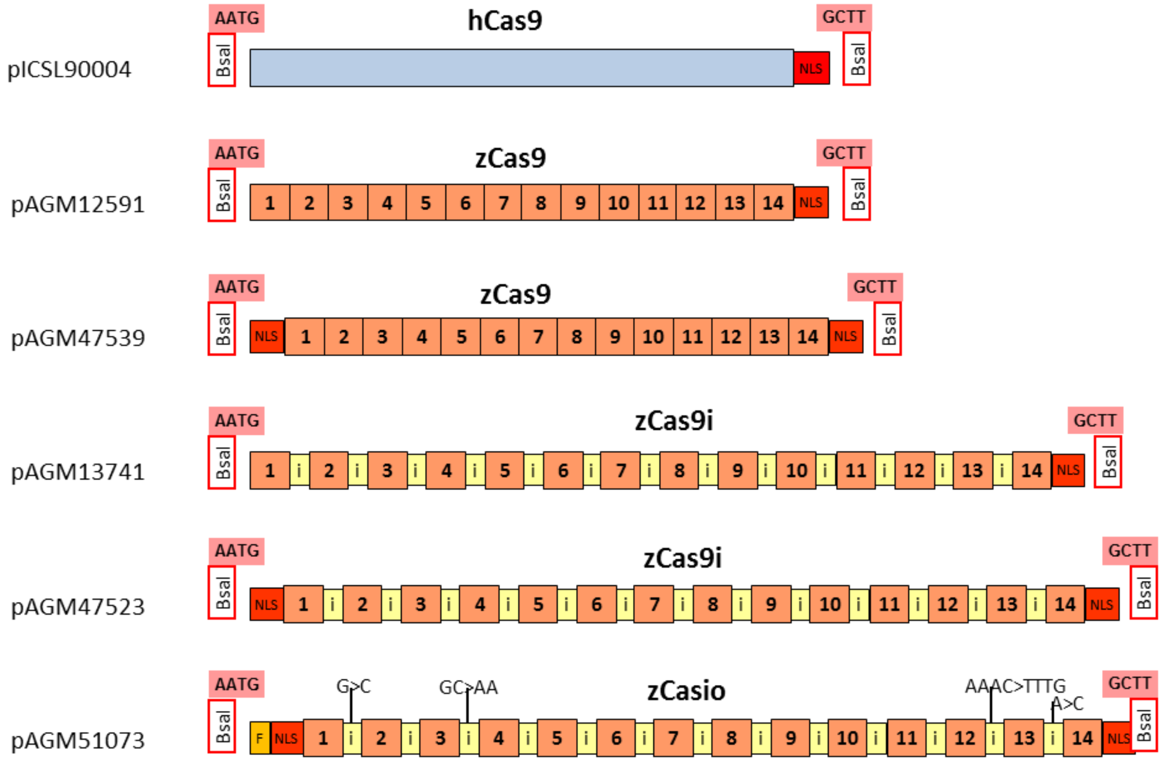

**Figure S2: Optimization of zCas9i to improve the predicted splicing efficiency.** Sequence changes were introduced into pAGM47523 to generate pAGM50631. Splice acceptor and donor sites predicted by Netgene2 are shown below the two Cas9 modules. A putative splice donor site present in pAGM47523 is not anymore predicted in pAGM50631. The confidence for the prediction of the splice sites circled in red are improved in pAGM50631. Note that in the end, the optimized sequence in pAGM50631 (zCas9io) was functional, but did not produce improved Cas9 activity in comparison to pAGM47523 (zCas9i) in Arabidopsis transformants.

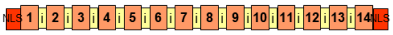


Donor splice sites, direct strand

| | pos 5'->3' | phase | strand | confidence | 5' exon intron 3' |
|---|---|---|---|---|---|
| | 322 | 1 | + | 1.00 | GATGGCGAAG^GTAAGGATTT H |
| Incorrect splice donor site | ~~342~~ | ~~2~~ | ~~+~~ | ~~0.74~~ | ~~TTATGATATA^GTATGCTTAT~~ |
| | 789 | 1 | + | 1.00 | AATGCATCAG^GTAACATTCC H |
| | 1114 | 1 | + | 0.95 | GCCCAAATAG^GTGCTCTTGA |
| | 1471 | 1 | + | 1.00 | ATTGATGGAG^GTAAGTTGTT H |
| | 1833 | 1 | + | 1.00 | TACTATGTAG^GTTAGTATCA H |
| | 2221 | 2 | + | 1.00 | TTCCTGTCAG^GTAAATCCTG H |
| | 2693 | 1 | + | 1.00 | AGATATACAG^GTAAGAGGTC H |
| | 3104 | 0 | + | 1.00 | GCTGGTGAAG^GTAAGTTCTG H |
| | 3572 | 0 | + | 1.00 | TGACAACAAG^GTAAAGCAAC H |
| | 3990 | 0 | + | 1.00 | GGAGGTAAAG^GTAAAGTTTC H |
| | 4472 | 1 | + | 1.00 | GGCGAAACAG^GTCTGTCTTT H |
| | 4941 | 1 | + | 1.00 | GAGGCTAAAG^GTAAAATATT H |
| | 5411 | 0 | + | 1.00 | CCTCGATAAG^GTAAGGACTT H |

Acceptor splice sites, direct strand

| pos 5'->3' | phase | strand | confidence | 5' intron exon 3' |
|---|---|---|---|---|
| 451 | 0 | + | 0.96 | GATTTTGCAG^GTTGACGACT |
| 879 | 1 | + | 1.00 | ACAATCTCAG^GTGTCGACGC H |
| 1221 | 1 | + | 0.95 | ACTTTTGTAG^GTGATCAGTA |
| 1571 | 1 | + | 1.00 | ATATTATTAG^GTGCAAGTCA H |
| 1953 | 1 | + | 1.00 | AATTCATTAG^GTCCACTCGC H |
| 2344 | 1 | + | 1.00 | GTTATAACAG^GTGAGCAGAA H |
| 2828 | 1 | + | 1.00 | TATGTTTTAG^GTTGGGGAAG H |
| 3253 | 0 | + | 1.00 | ATTGCTACAG^GTCATGGGCC H |
| 3692 | 0 | + | 0.95 | CGTGTTATAG^GTCCTTACGC |
| 4125 | 0 | + | 0.95 | GGTTACGCAG^GTGATCACCC |
| 4592 | 1 | + | 1.00 | TTTCTTGCAG^GTGAGATTGT H |
| 5099 | 1 | + | 0.87 | TACAAACCAG^GTTACAAAGA |
| 5519 | 0 | + | 0.94 | ACAATTAAAG^GTGCTTTCCG |

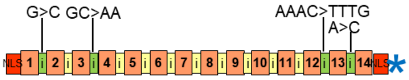


Donor splice sites, direct strand

| pos 5'->3' | phase | strand | confidence | 5' exon intron 3' |
|---|---|---|---|---|
| 322 | 0 | + | 1.00 | GATGGCGAAG^GTAAGGATTT H |
| 789 | 1 | + | 1.00 | AATGCATCAG^GTAACATTCC H |
| 1114 | 1 | + | 1.00 | GCCCAAATAG^GTAATCTTGA H |
| 1471 | 1 | + | 1.00 | ATTGATGGAG^GTAAGTTGTT H |
| 1833 | 1 | + | 1.00 | TACTATGTAG^GTTAGTATCA H |
| 2221 | 2 | + | 1.00 | TTCCTGTCAG^GTAAATCCTG H |
| 2693 | 1 | + | 1.00 | AGATATACAG^GTAAGAGGTC H |
| 3104 | 0 | + | 1.00 | GCTGGTGAAG^GTAAGTTCTG H |
| 3572 | 0 | + | 1.00 | TGACAACAAG^GTAAAGCAAC H |
| 3990 | 0 | + | 1.00 | GGAGGTAAAG^GTAAAGTTTC H |
| 4472 | 1 | + | 1.00 | GGCGAAACAG^GTCTGTCTTT H |
| 4941 | 1 | + | 1.00 | GAGGCTAAAG^GTAAAATATT H |
| 5411 | 0 | + | 1.00 | CCTCGATAAG^GTAAGGACTT H |

Acceptor splice sites, direct strand

| pos 5'->3' | phase | strand | confidence | 5' intron exon 3' |
|---|---|---|---|---|
| 451 | 0 | + | 0.96 | GATTTTGCAG^GTTGACGACT |
| 879 | 1 | + | 1.00 | ACAATCTCAG^GTGTCGACGC H |
| 1221 | 1 | + | 0.95 | ACTTTTGTAG^GTGATCAGTA |
| 1571 | 1 | + | 1.00 | ATATTATTAG^GTGCAAGTCA H |
| 1953 | 1 | + | 1.00 | AATTCATTAG^GTCCACTCGC H |
| 2344 | 1 | + | 1.00 | GTTATAACAG^GTGAGCAGAA H |
| 2828 | 1 | + | 1.00 | TATGTTTTAG^GTTGGGGAAG H |
| 3253 | 0 | + | 1.00 | ATTGCTACAG^GTCATGGGCC H |
| 3692 | 0 | + | 0.95 | CGTGTTATAG^GTCCTTACGC |
| 4125 | 0 | + | 0.95 | GGTTACGCAG^GTGATCACCC |
| 4592 | 1 | + | 1.00 | TTTCTTGCAG^GTGAGATTGT H |
| 5099 | 1 | + | 1.00 | TACTTTGCAG^GTTACAAAGA |
| 5519 | 0 | + | 1.00 | ACAATTACAG^GTGCTTTCCG H |

**Figure S3: Pictures of primary transformants transformed with constructs targeting *CPC* and *TRY***

pAGM51511

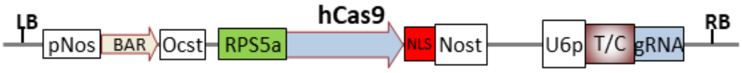


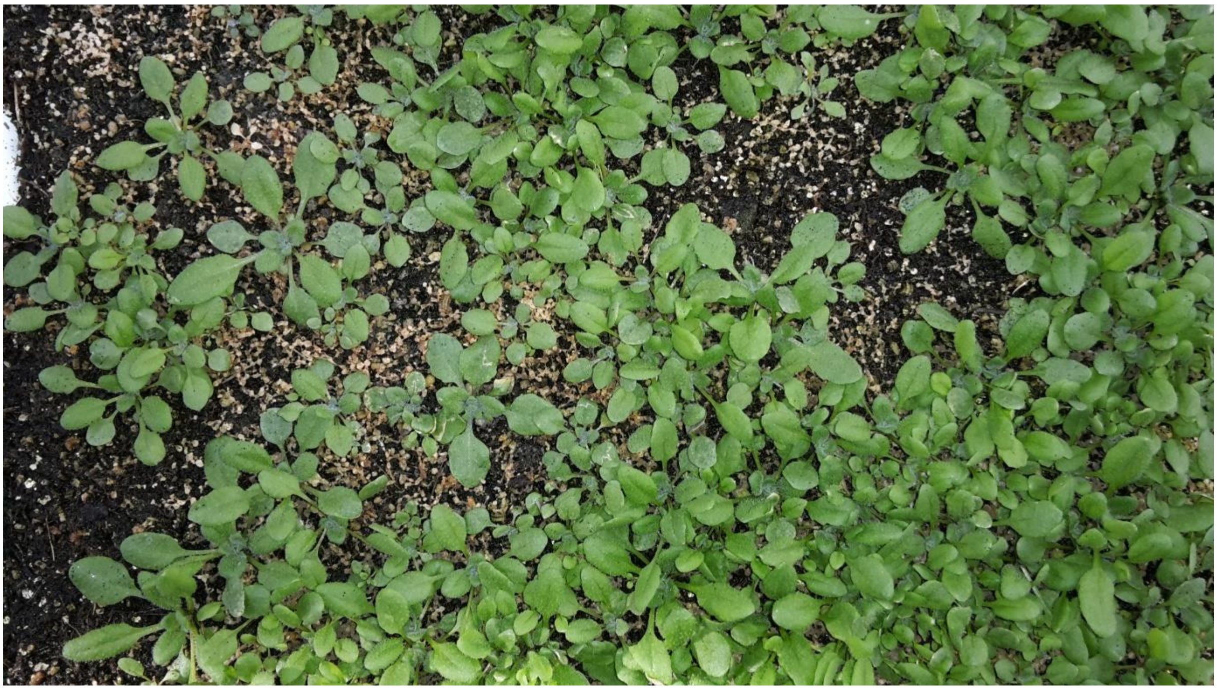

pAGM51613

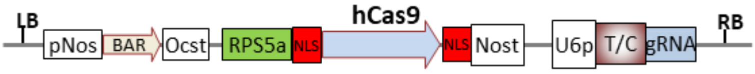


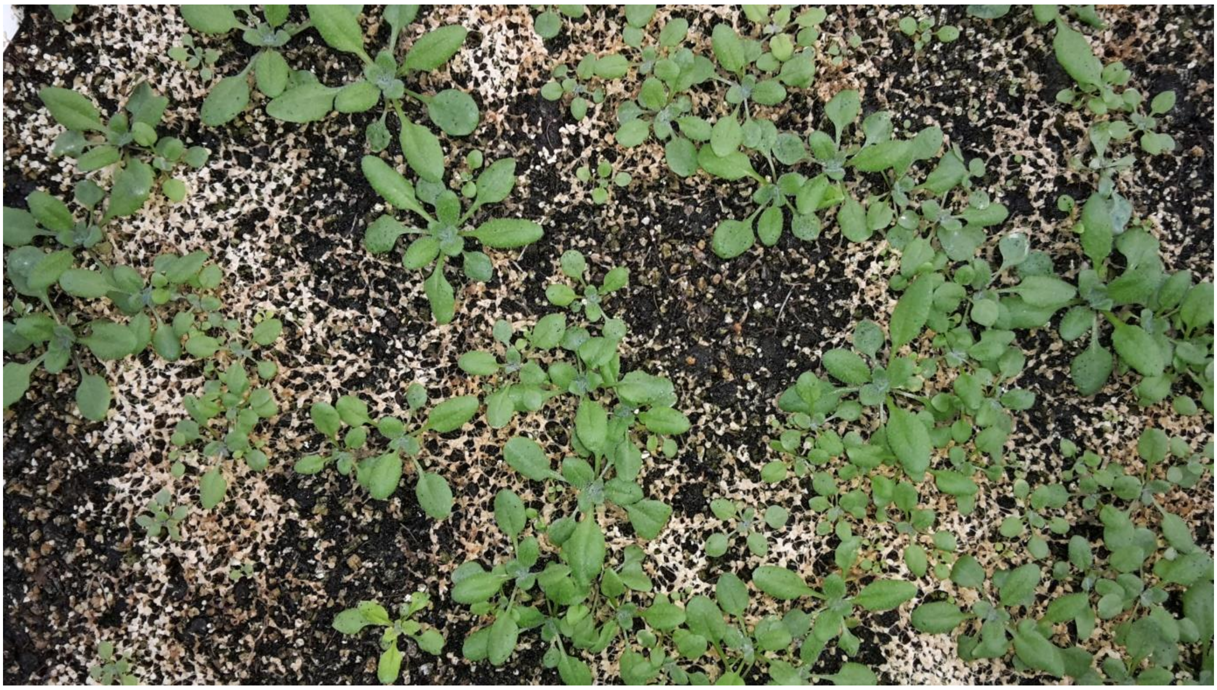

**Figure S3** ***(continued)***

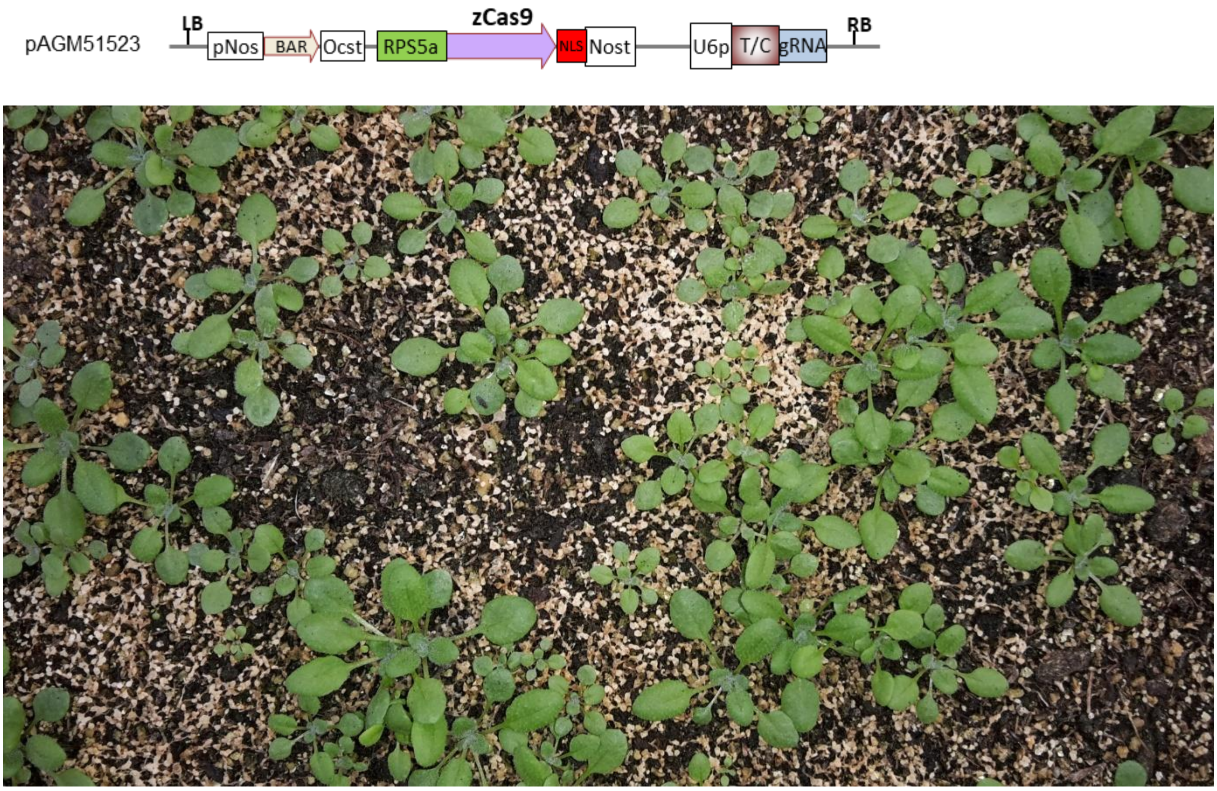


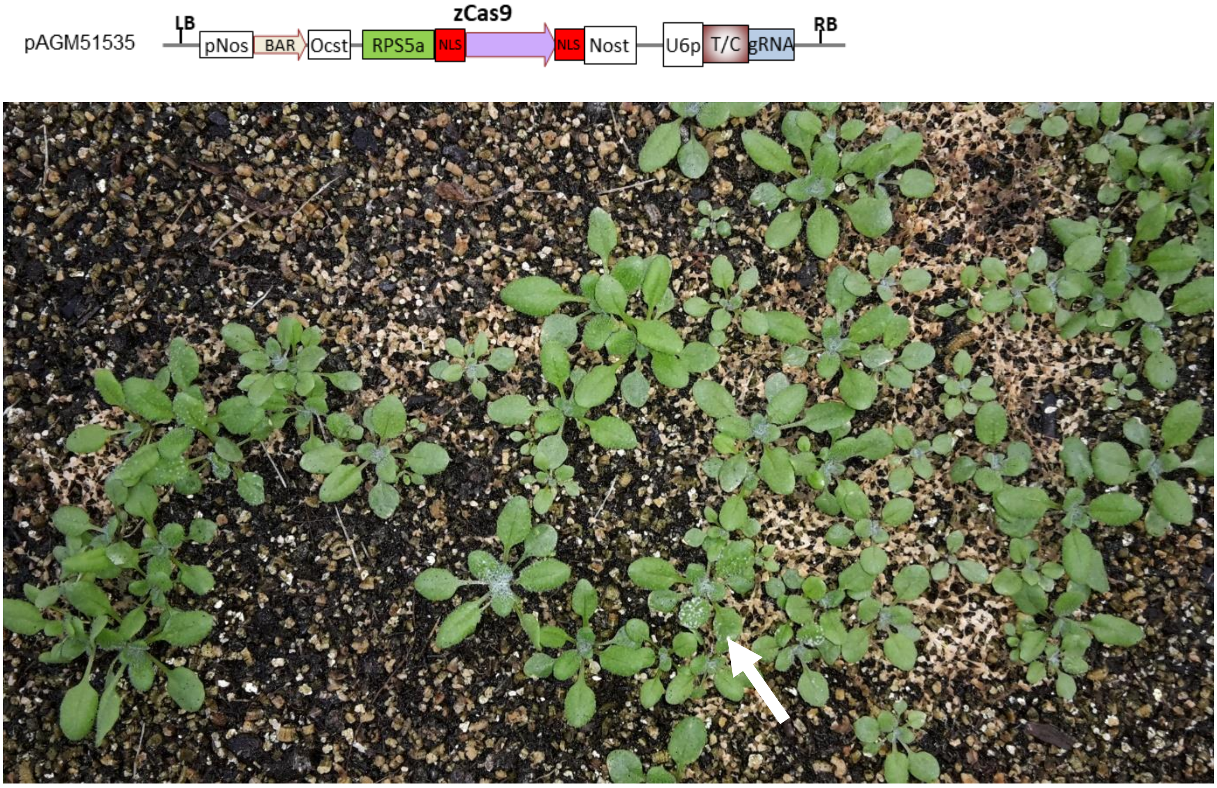

**Figure S3** ***(continued)***

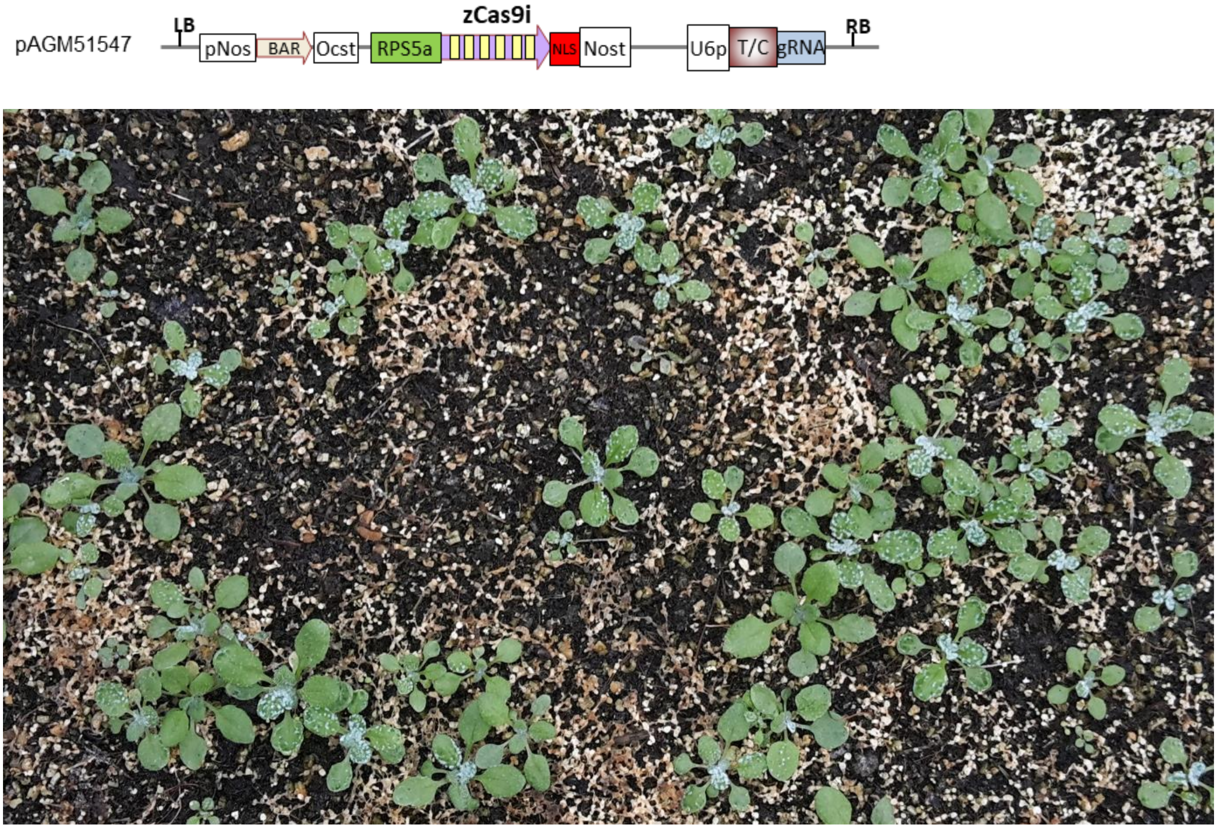


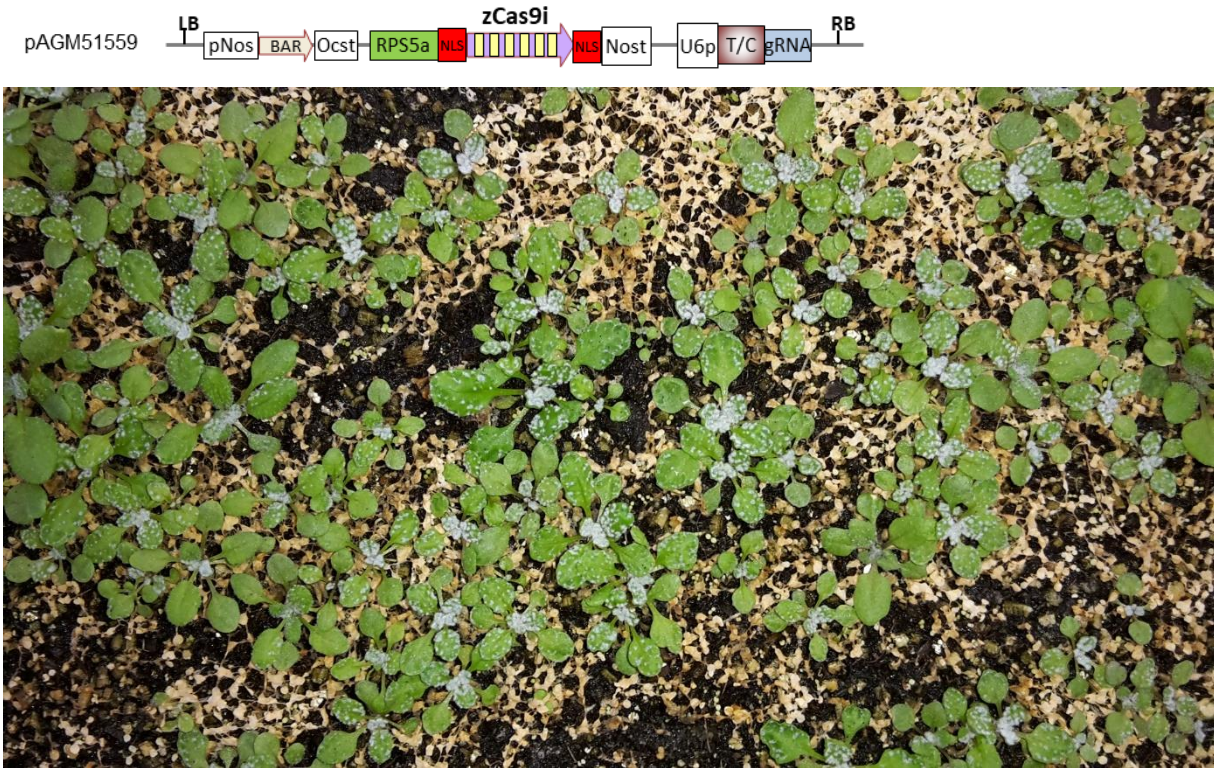

**Figure S3** ***(continued)***

pAGM51561

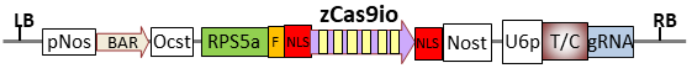


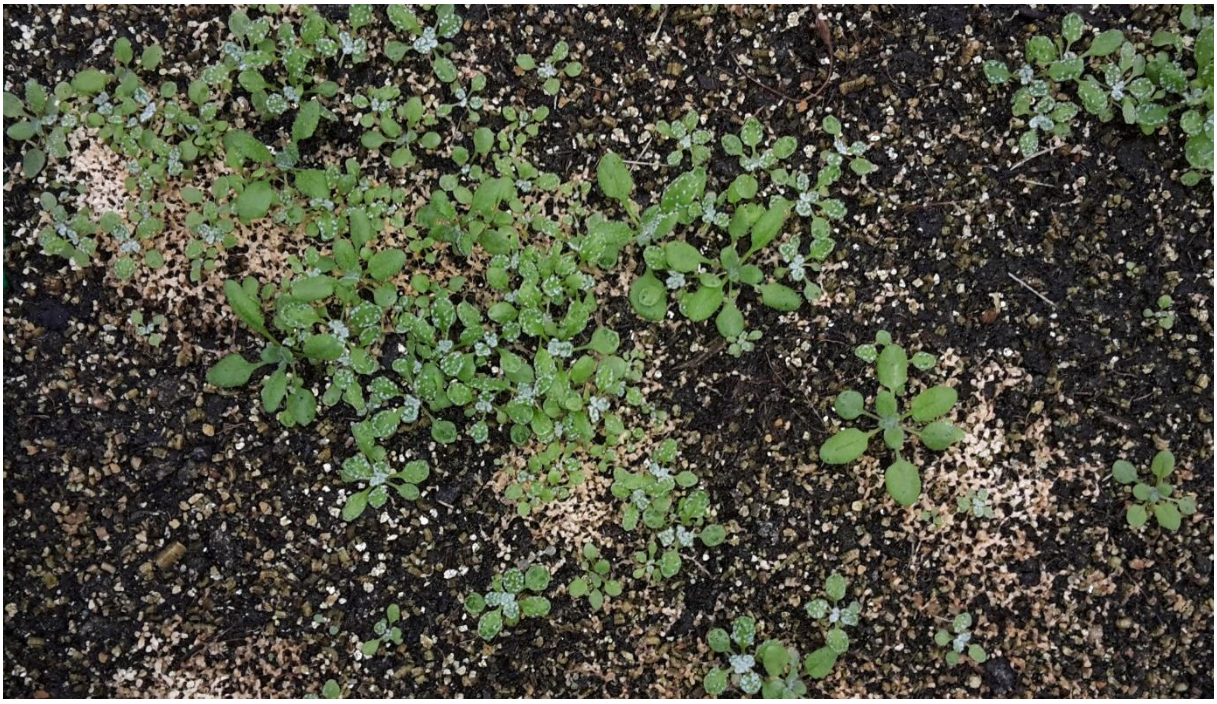

pAGM52451
(low copy)

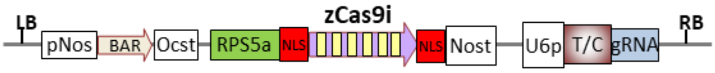


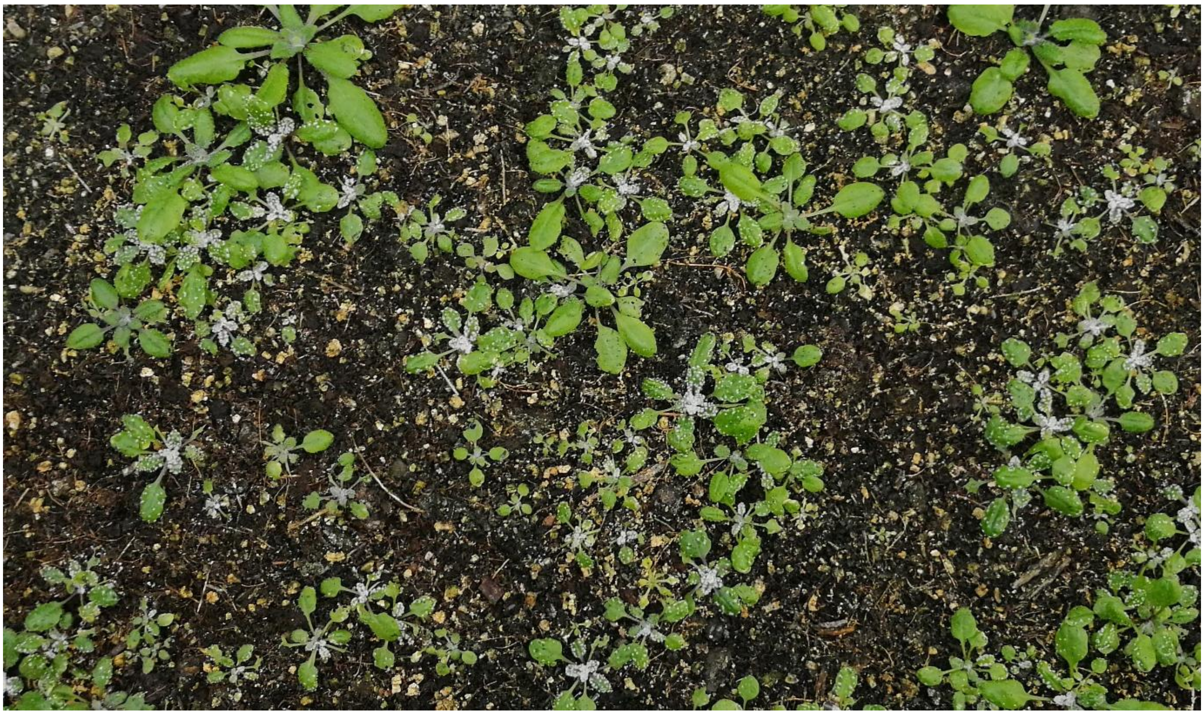

**Figure S4: Analysis of Arabidopsis transformants for presence of a T-DNA.** The genomic DNA of Arabidopsis primary transformants obtained by transformation of pAGM51559 and pAGM51561 was analyzed with two primer pairs located at the beginning of the Cas9 coding sequence and in the region of the guide RNA (two regions of the T-DNA separated by more than 5 kb). Plants with either a mutant or WT phenotype were analyzed, but chimeric plants displaying a mutant phenotype in some leafs but not others were also analyzed, both in sectors with the WT phenotype and in sectors with the mutant phenotype. Arabidopsis WT DNA was used as a negative control. The presence of two bands of the expected sizes suggests the presence of a complete T-DNA in most plants. Some plants have only one of the 2 bands (pAGM51561 transformant 25) indicating the presence of an incomplete T-DNA. Plant pAGM51559 transformant 8 only shows a smear as in the negative control with WT DNA, suggesting that this plant probably lacks most of the T-DNA r.

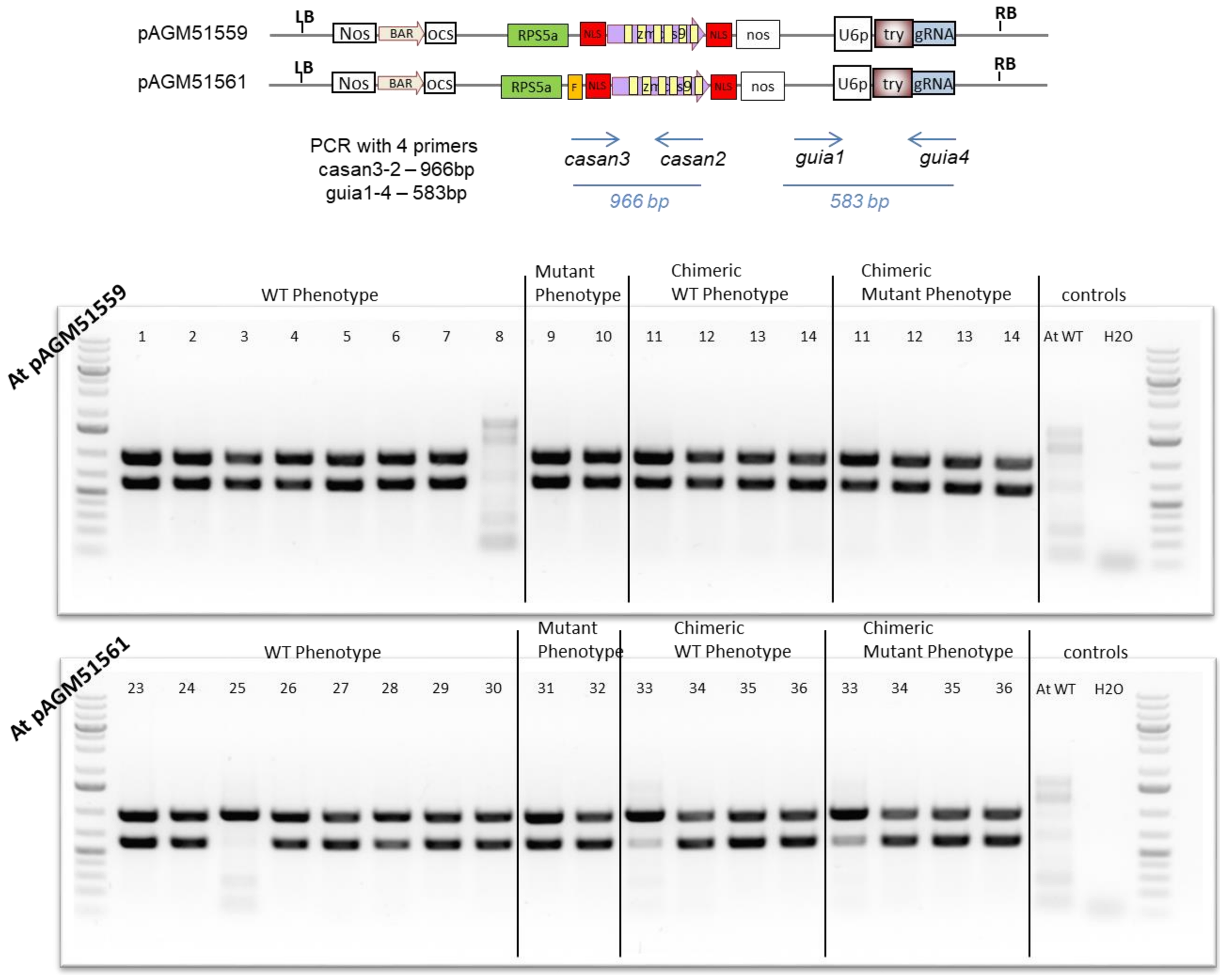

**Figure S5a: Sequence analysis of *TRY* and *CPC* genes in the pAGM51559 transformants.** PCR was performed with primers cpcan1 (tt ggtctc a ACAT gtcagaactcactttggctagtttgg) and cpcan2 (tt ggtctc a ACAA atcatgtgtcgatggaggctgg) for the *CPC* gene and sequenced with primer cpcan1. PCR was performed with primers tryan1 (tt ggtctc a ACAT ggggaagcacatggtgtccac) and tryan2 (tt ggtctc a ACAA gttgtggatataaaagtcgtagacgag) for *TRY* and sequenced with primer tryan2. WT1 to WT8 are 8 pAGM51559 transformants with WT phenotype. Mut9 and Mut10 are two plants with full knock out phenotype. * refers to PCR products that were later cloned for the *CPC* gene, and 10 resulting clones sequenced (see Figure S6). The sequences traces of plants with no mutations are not shown, and "WT" is written instead. A WT sequence trace is shown at the bottom of the figure.

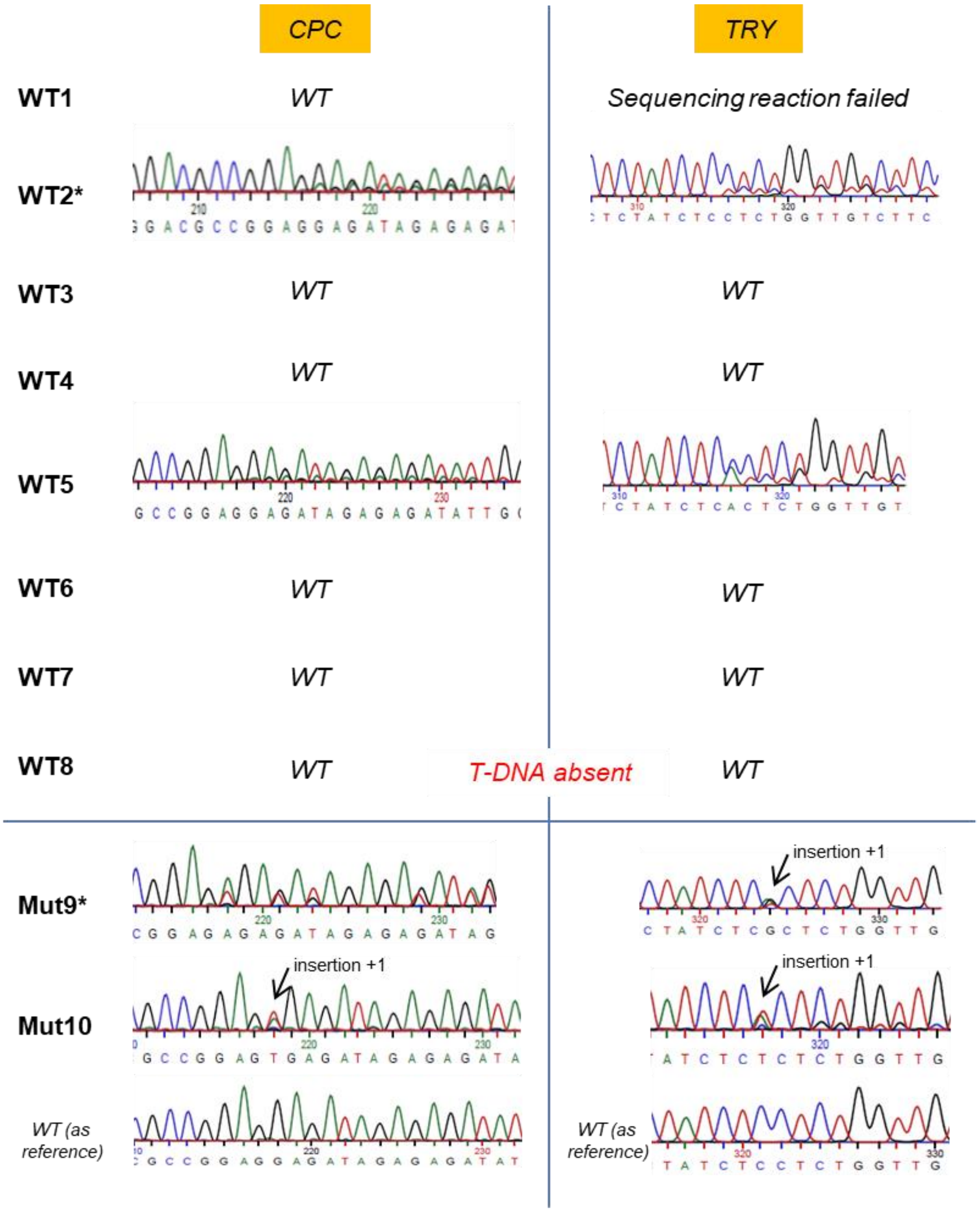

**Figure S5b: Sequence analysis of *TRY* and *CPC* genes in the pAGM51559 transformants**. PCR was performed with primers tryan1 (tt ggtctc a ACAT ggggaagcacatggtgtccac) and tryan2 (tt ggtctc a ACAA gttgtggatataaaagtcgtagacgag) and sequenced with primer tryan2. CWT11 to 14 and cmut 11 to 14 are DNA samples extracted from 4 pAGM51559 transformants with chimeric mutant/WT phenotype. DNA was extracted with leaves with a WT phenotype in samples CWT11 to 14, and from leaves with a mutant phenotype in Cmut11 to 14. * refers to PCR products that were later cloned and 10 resulting clones sequenced (see Figure S6).

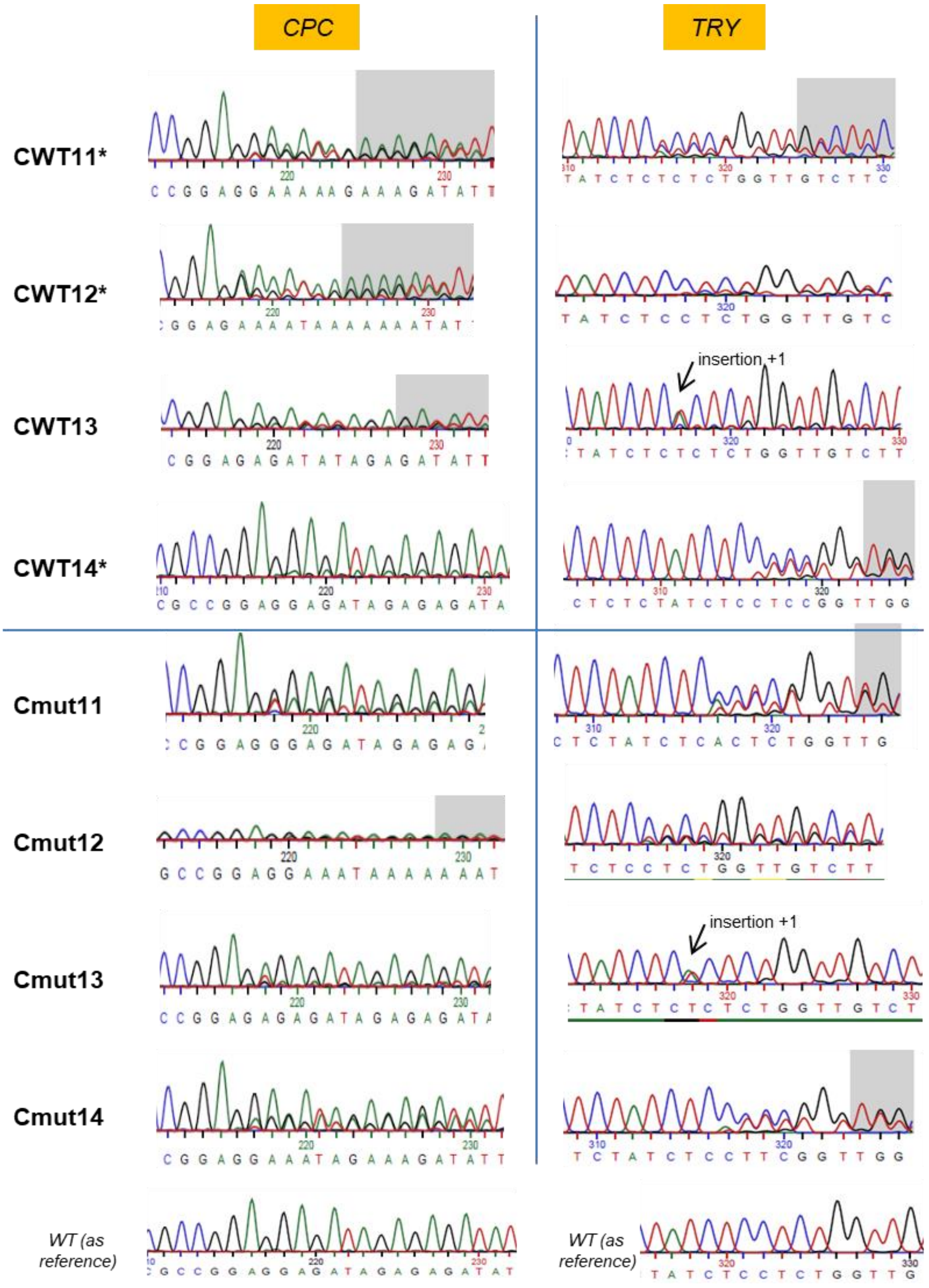

**Figure S6: Sequences of cloned PCR products from pAGM51559 transformants.** PCR products amplified from the *CPC* gene from Figure S5 were cloned in vector pAGM1311 using BsaI and ligase. 10 of the resulting clones were sequenced using vector primer Moclofwd (agcgaggaagcggaagagcg). The sequence alignment of the successful sequencing reactions is shown. Sequence annotated PCR cpcan1-2 corresponds to the WT sequence.

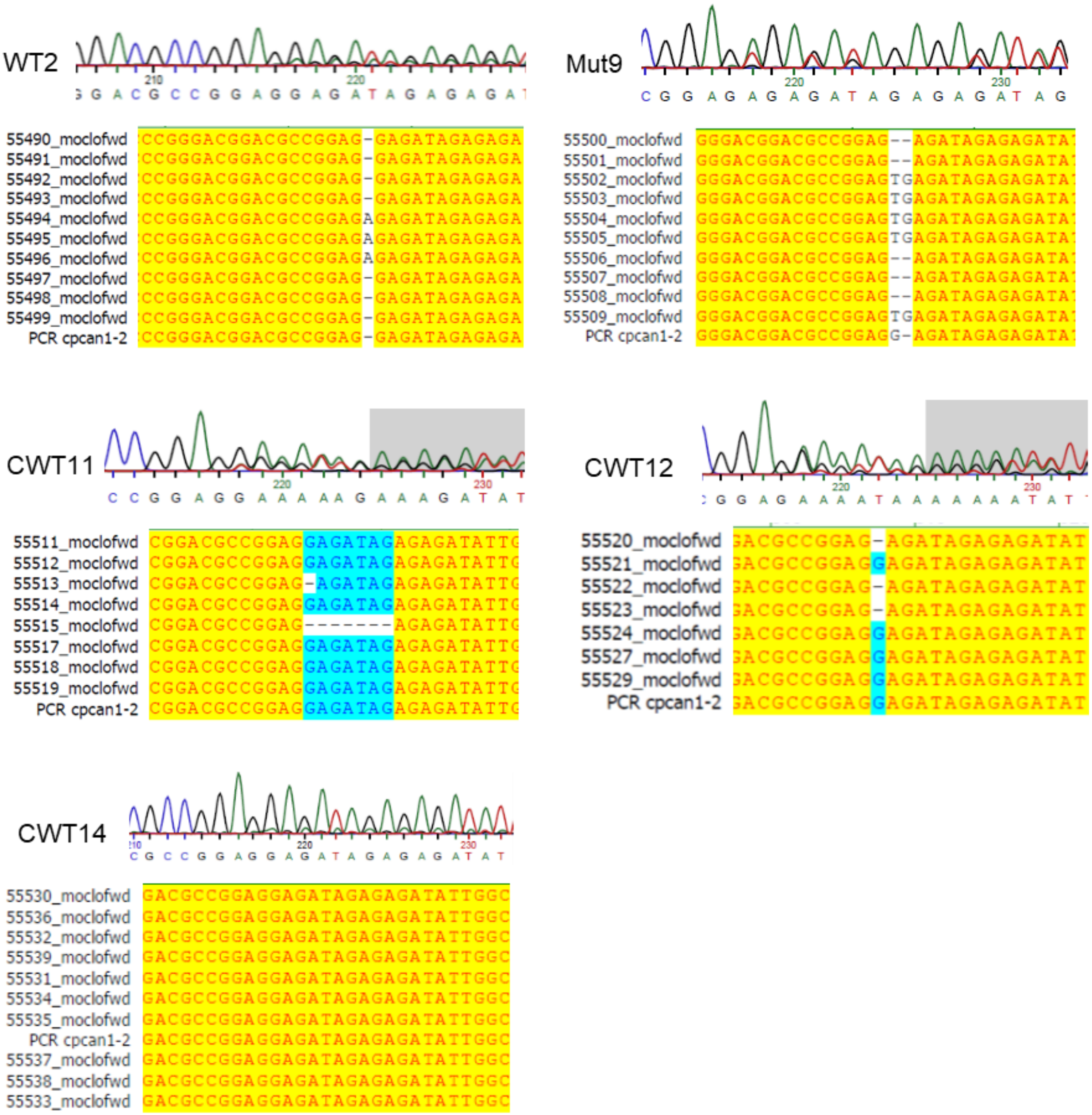

**Figure S7: Sequence analysis of *CPC* in pAGM51561 transformants.** PCR was performed with primers cpcan1 (tt ggtctc a ACAT gtcagaactcactttggctagtttgg) and cpcan2 (tt ggtctc a ACAA atcatgtgtcgatggaggctgg) for the *CPC* gene and sequenced with primer cpcan1. WT23 to WT30 are 8 pAGM51561 transformants with WT phenotype. Mut31 and Mut32 are two plants with full knock out phenotype. CWT33 to 36 and cmut33 to 36 are DNA samples extracted from 4 pAGM51559 transformants with chimeric mutant/WT phenotype. DNA was extracted from leaves with a WT phenotype in samples CWT33 to 36, and from leaves with a mutant phenotype in Cmut33 to 36. * refers to PCR products that were later cloned for the *CPC* gene, and 10 resulting clones sequenced (see Figure S8). The sequence traces of plants with no mutations are not shown, and "WT" is written instead.

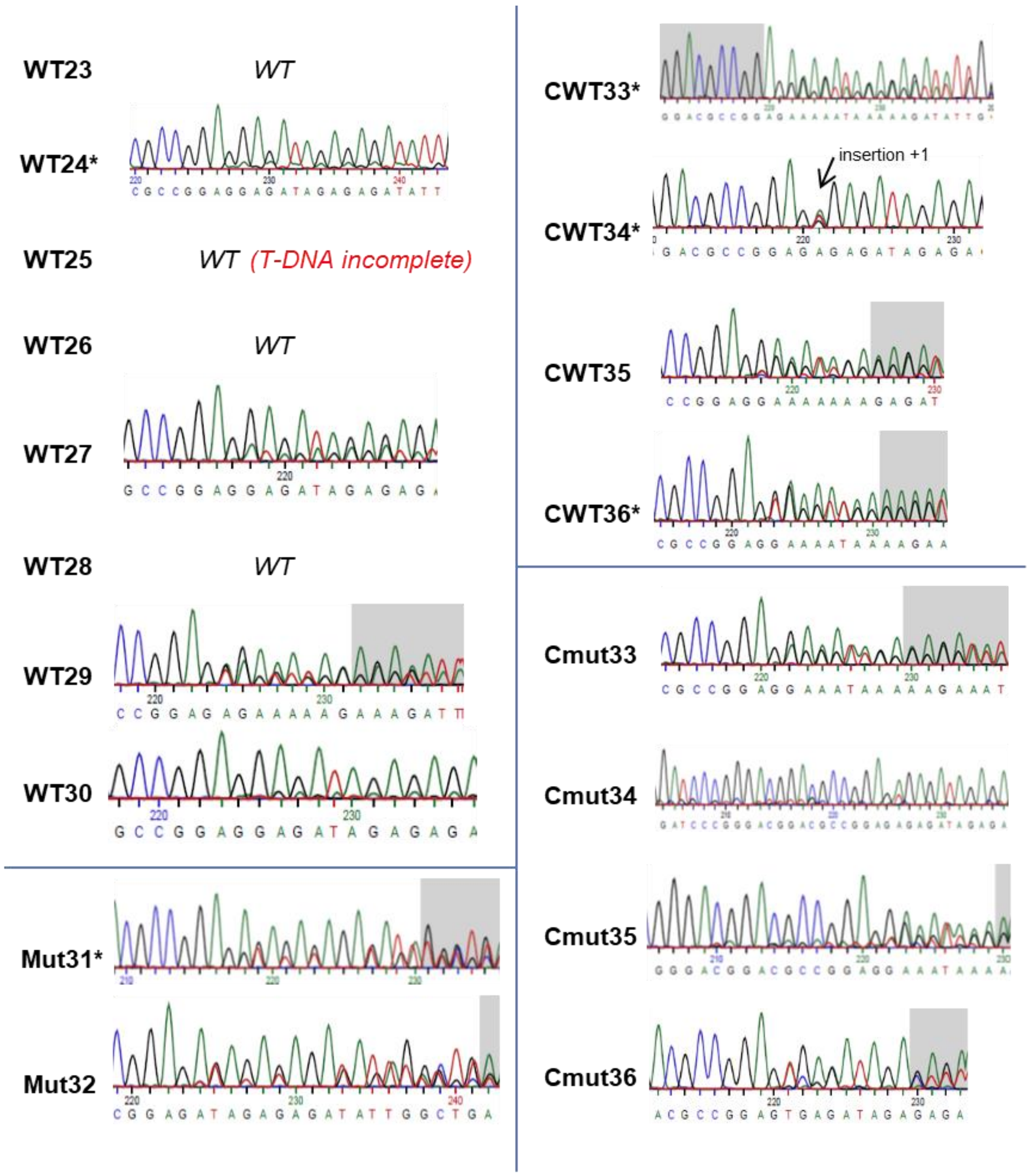

**Figure S8: Sequences analysis of cloned PCR products from pAGM51561 transformants.** PCR products from the *CPC* gene from Fig S7 were cloned in vector pAGM1311 using BsaI and ligase. 10 of the resulting clones were sequenced using vector primer Moclofwd (agcgaggaagcggaagagcg). The sequence alignment of the successful sequencing reactions is shown. Sequence annotated PCR cpcan1-2 corresponds to the WT sequence.

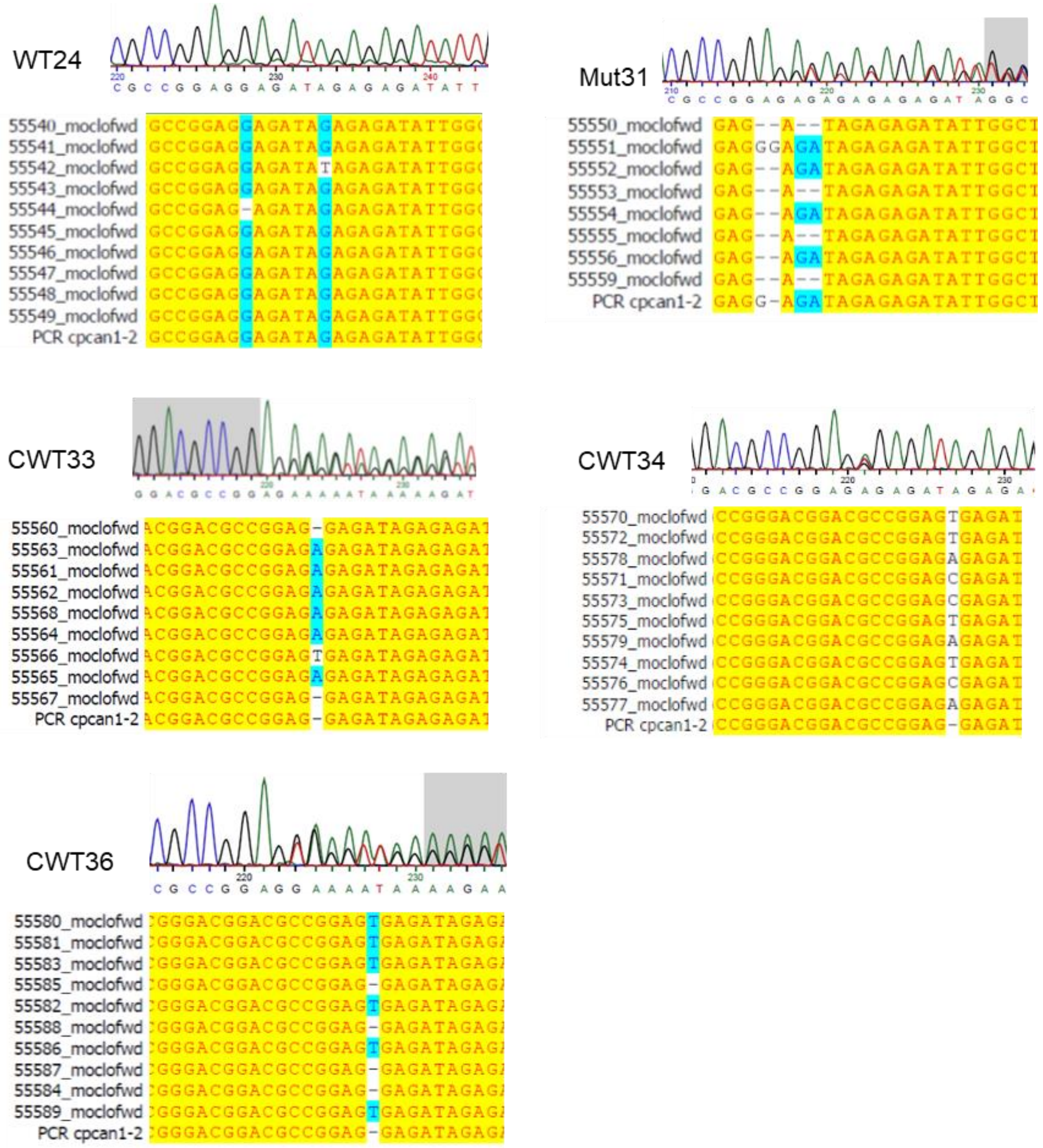

**Figure S9: Vectors for cloning Cas9 constructs.** (a) Vectors for cloning single guide RNAs. Two BsaI sites located between the Arabidopsis U6 promoter and the invariable region of the guide RNA allow cloning of a guide RNA by ligation of two 24 nt oligonucleotides. A FAST cassette is present between the guide RNA cassette and the right border for transgene counter-selection by seed fluorescence. (b) Vectors for cloning 1 to 6 guide RNAs using the MoClo system (see Figure S16).

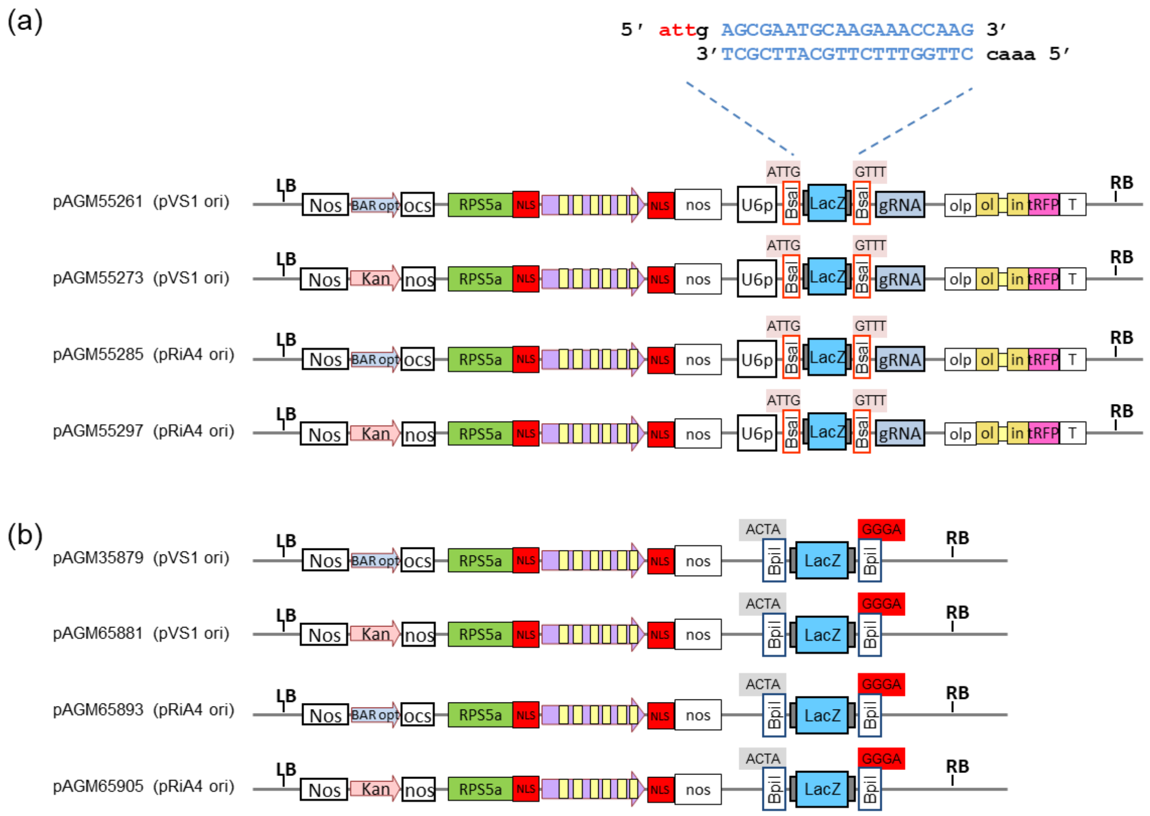

**Figure S10: Cas9 mutagenesis in *Nicotiana benthamiana*, overview of results. (a)** Constructs were made for site-targeted mutagenesis in *Nicotiana benthamiana* using hCas9 and zCas9i (only for target sites rt1 and rt2) or zCas9i only for all other target sites. The number of plants displaying mutations as a ratio of all tested plants is shown. Note that not all transformants were analyzed for presence of the T-DNA**. (b)** Sequence of the target sites; PAM sequence underlined.

(a)

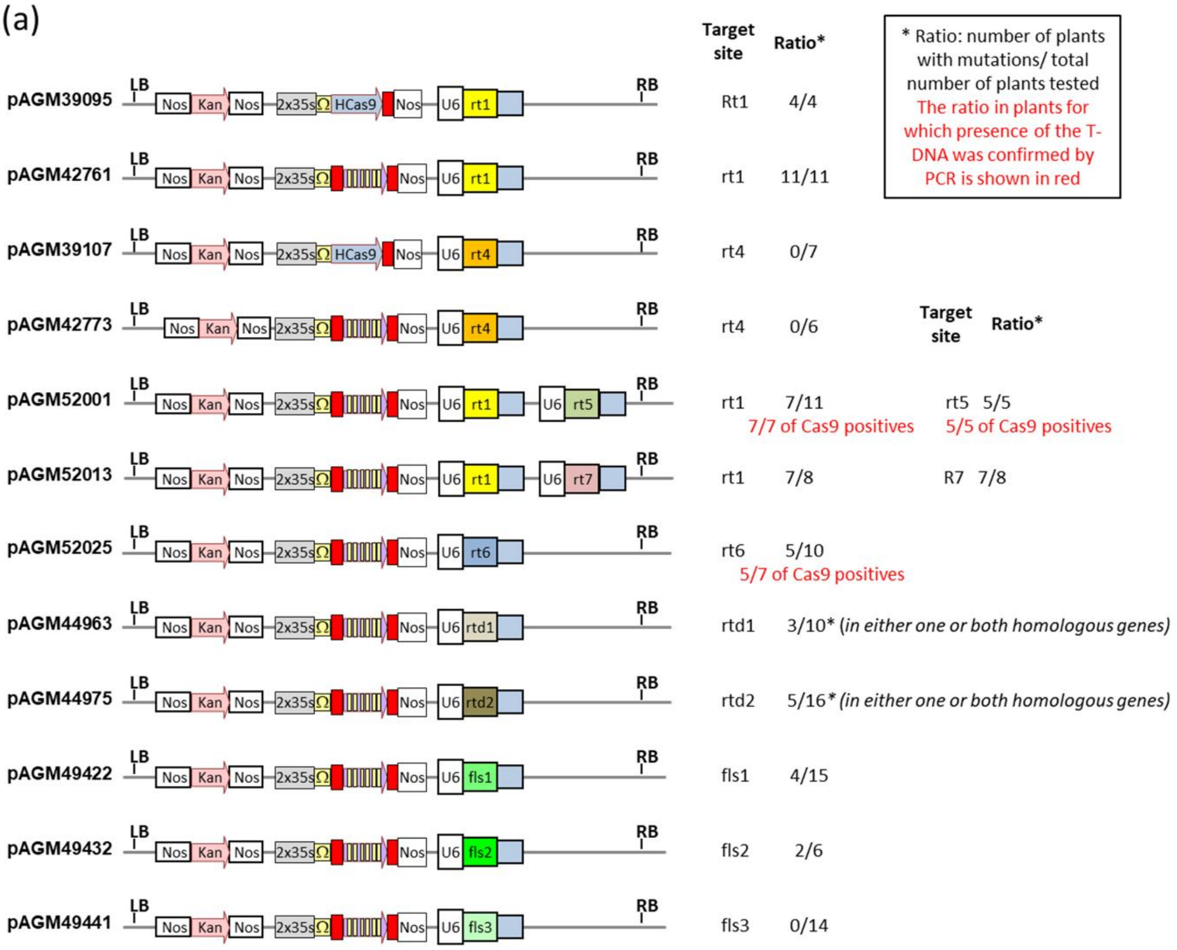


(b)

| | |
|---|---|
| rt1 | agggggactatgtgagtcgt ggg |
| rt4 | aaatctactctggcaagcat tgg |
| rt5 | gagggggactatgtgagtcg tgg |
| rt6 | caagacttgttctcaaatgg agg |
| rt7 | gctgtgctttccgaaccagg agg |
| rtd1 | ggaggtagaccttcaacttg agg |
| rtd2 | gtgagtagagtcttgatttg tgg |
| fls1 | gagatgattgcaaagaatcc agg |
| fls2 | gttagaagcccatgaaatga tgg |
| fls3 | gaaatgatggaggcagctgg tgg |

**Figure S11: Analysis of mutations in *Catharanthus roseus* obtained with construct pSB312**. pSB312 contains 8 guide RNAs targeting 3 genes (*JAM2*, *JAM3*, and *RMT1*)

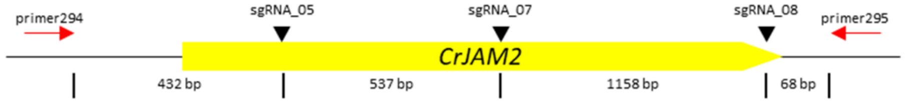


| Line | Allele | sgRNA_05 | sgRNA_07 | sgRNA_08 |
|---|---|---|---|---|
| wt gDNA/rSB02_310 lines | A (homozygous) | | | |
| rSB02_312_03_2 | A | **550 bp deletion** | | **-4-4-1-2** |
| | B | **1711 bp deletion** | | |
| rSB02_312_18_1 | A (homozygous) | -3 | **-2** | **-1** |
| rSB02_312_23_5 | A | **-5** | **+1** | **+1** |
| | B | **558 bp deletion** | | -16 & T to G |
| rSB02_312_26_1 | A | **+1** | **-14** | **-19** |
| | B | **reverse complementary & additional ~350 bp deletion** | | **-34** |

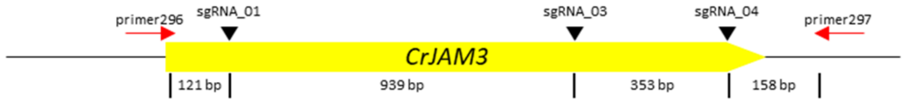


| Line | Allele | sgRNA_01 | sgRNA_03 | sgRNA_04 |
|---|---|---|---|---|
| wt gDNA/rSB02_310 lines | A (homozygous) | | | |
| rSB02_312_03_2 | A | **reverse complementary** | | -3 |
| | B | -15 | **-5** | -63 |
| rSB02_312_18_1 | A (homozygous) | **reverse complementary (sgRNA_01-03) & 344 bp deletion (sgRNA_03-04)** | | |
| rSB02_312_23_5 | A (homozygous) | **1298 bp deletion** | | |
| rSB02_312_26_1 | A | **-64** | **+1** | **-28** |
| | B | **-1** | **+1** | **-28** |

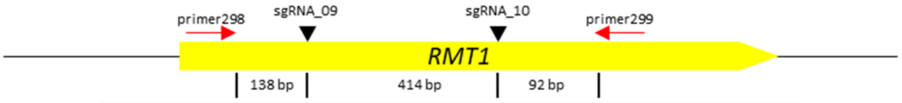


| Line | Allele | sgRNA_09 | sgRNA_10 |
|---|---|---|---|
| wt gDNA/rSB02_310 lines | A (homozygous) | | |
| rSB02_312_03_2 | A | **1kb insertion from 3' of RMT1** | -3 |
| | B | **+1-8** | -3 |
| rSB02_312_18_1 | A | **-10** | **-25** |
| | B | -27 | -72+9 |
| rSB02_312_23_5 | A | -27 | **-1** |
| | B | -27 | **-13** |
| rSB02_312_26_1 | A | -27 | **+1** |
| | B | -27 | **-8** |

**Figure S12:** Nucleotide sequences of the Cas9 and NLS coding sequences

Human codon-optimized Cas9-NLS (hCas9) from pAGM51511 and pAGM51613 (NLS in red)

ATGgacaagaagtactccattgggctcgatatcggcacaaacagcgtcggctgggccgtcattacggacgagtacaaggtgccgagcaaaaa
attcaaagttctgggcaataccgatcgccacagcataaagaagaacctcattggcgccctcctgttcgactccggggagacggccgaagccacg
cggctcaaaagaacagcacggcgcagatatacccgcagaaagaatcggatctgctacctgcaggagatctttagtaatgagatggctaaggtgg
atgactctttcttccataggctggaggagtcctttttggtggaggaggataaaaagcacgagcgccacccaatctttggcaatatcgtggacgagg
tggcgtaccatgaaaagtacccaaccatatatcatctgaggaagaagcttgtagacagtactgataaggctgacttgcggttgatctatctcgcgc
tggcgcatatgatcaaatttcggggacacttcctcatcgaggggggacctgaacccagacaacagcgatgtcgacaaactctttatccaactggttc
agacttacaatcagcttttcgaagagaacccgatcaacgcatccggagttgacgccaaagcaatcctgagcgctaggctgtccaaatcccggcg
gctcgaaaacctcatcgcacagctccctggggagaagaagaacggcctgtttggtaatcttatcgccctgtcactcgggctgacccccaactttaa
atctaacttcgacctggccgaagatgccaagcttcaactgagcaaagacacctacgatgatgatctcgacaatctgctggcccagatcggcgacc
agtacgcagacctttttttggcggcaaagaacctgtcagacgccattctgctgagtgatattctgcgagtgaacacggagatcaccaaagctccgc
tgagcgctagtatgatcaagcgctatgatgagcaccaccaagacttgactttgctgaaggcccttgtcagacagcaactgcctgagaagtacaag
gaaattttcttcgatcagtctaaaaatggctacgccggatacattgacggcggagcaagccaggaggaattttacaaatttattaagcccatcttg
gaaaaaatggacggcaccgaggagctgctggtaaagcttaacagagaagatctgttgcgcaaacagcgcactttcgacaatggaagcatcccc
caccagattcacctgggcgaactgcacgctatcctcaggcggcaagaggatttctacccctttttgaaagataacagggaaaagattgagaaaat
cctcacatttcggataccctactatgtaggcccccctcgcccggggaaattccagattcgcgtggatgactcgcaaatcagaagagactatcactcc
ctggaacttcgaggaagtcgtggataagggggcctctgcccagtccttcatcgaaaggatgactaactttgataaaaatctgcctaacgaaaagg
tgcttcctaaacactctctgctgtacgagtacttcacagtttataacgagctcaccaaggtcaaatacgtcacagaagggatgagaaagccagca
ttcctgtctggagagcagaagaaagctatcgtggacctcctcttcaagacgaaccggaaagttaccgtgaaacagctcaaagaagattatttcaa
aaagattgaatgtttcgactctgttgaaatcagcggagtggaggatcgcttcaacgcatccctgggaacgtatcacgatctcctgaaaatcattaa
agacaaggacttcctggacaatgaggagaacgaggacattcttgaggacattgtcctcacccttacgttgtttgaagatagggagatgattgaag
aacgcttgaaaacttacgctcatctcttcgacgacaaagtcatgaaacagctcaagaggcgccgatatacaggatgggggcggctgtcaagaaa
actgatcaatgggatccgagacaagcagagtggaaagacaatcctggattttcttaagtccgatggatttgccaaccggaacttcatgcagttga
tccatgatgactctctcacctttaaggaggacatccagaaagcacaagtttctggccagggggacagtctccacgagcacatcgctaatcttgca
ggtagcccagctatcaaaaagggaatactgcagaccgttaaggtcgtggatgaactcgtcaaagtaatgggaaggcataagcccgagaatatc
gttatcgagatggcccgagagaaccaaactacccagaagggacagaagaacagtagggaaaggatgaagaggattgaagagggtataaaag
aactggggtcccaaatccttaaggaacacccagttgaaaacacccagcttcagaatgagaagctctacctgtactacctgcagaacggcaggga
catgtacgtggatcaggaactggacatcaatcggctctccgactacgacgtggatcatatcgtgccccagtcttttctcaaagatgattctattgat
aataaagtgttgacaagatccgataaaaatagagggaagagtgataacgtcccctcagaagaagttgtcaagaaaatgaaaaattattggcgg
cagctgctgaacgccaaactgatcacacaacggaagttcgataatctgactaaggctgaacgaggtggcctgtctgagttggataaagccggctt
catcaaaaggcagcttgttgagacacgccagatcaccaagcacgtggcccaaattctcgattcacgcatgaacaccaagtacgatgaaaatgac
aaactgattcgagaggtgaaagttattactctgaagtctaagctggtttcagatttcagaaaggactttcagttttataaggtgagagagatcaac
aattaccaccatgcgcatgatgcctacctgaatgcagtggtaggcactgcacttatcaaaaaatatcccaagcttgaatctgaatttgtttacgga
gactataaagtgtacgatgttaggaaaatgatcgcaaagtctgagcaggaaataggcaaggccaccgctaagtacttcttttacagcaatattat
gaatttttcaagaccgagattacactggccaatggagagattcggaagcgaccacttatcgaaacaaacggagaaacaggagaaatcgtgtg
ggacaagggtagggatttcgcgacagtccggaaggtcctgtccatgccgcaggtgaacatcgttaaaaagaccgaagtacagaccggaggctt
ctccaaggaaagtatcctcccgaaaaggaacagcgacaagctgatcgcacgcaaaaaagattgggaccccaagaaatacggcggattcgattc
tcctacagtcgcttacagtgtactggttgtggccaaagtggagaaagggaagtctaaaaaactcaaaagcgtcaaggaactgctgggcatcaca
atcatggagcgatcaagcttcgaaaaaaaccccatcgactttctcgaggcgaaaggatataaagaggtcaaaaaagacctcatcattaagcttc
ccaagtactctctctttgagcttgaaaacggccggaaacgaatgctcgctagtgcgggcgagctgcagaaaggtaacgagctggcactgccctct
aaatacgttaatttcttgtatctggccagccactatgaaaagctcaaaggatctcccgaagataatgagcagaagcagctgttcgtggaacaaca
caaacactaccttgatgagatcatcgagcaaataagcgaattctccaaaagagtgatcctcgccgacgctaacctcgataaggtgctttctgctta
caataagcacagggataagcccatcagggagcaggcagaaaacattatccacttgtttactctgaccaacttgggcgcgcctgcagccttcaagt

acttcgacaccaccatagacagaaagcggtacacctctacaaaggaggtcctggacgccacactgattcatcagtcaattacggggctctatgaa
acaagaatcgacctctctcagctcggtggagac **agcagggctgaccccaagaagaagaggaaggtg** TGA

Cas9 N-terminal NLS of pAGM51563 and pAGM51535

**ATGgcttctagcccaccgaagaagaagcggaaggtcagctggaaa**

*Zea mays* codon-optimized Cas9-NLS (zCas9) of pAGM51523 and pAGM51535 (NLS in red)

atggacaagaagtacagcattggacttgatattggtacgaactcagttgggtgggccgttatcaccgatgaatacaaggtaccttcgaagaaatt
taaagtgctgggcaacacagataggcacagcattaagaagaacttgatcggagctctgctctttgactctggagaaaccgcggaggcgacaagg
cttaaacgtactgcgaggagaaggtacactcgcaggaagaacagaatctgttatctccaagagatctttagcaacgagatggcgaaggttgacg
actcgttcttccatcgcctcgaggaatccttcctggtagaggaagataagaaacacgagcgtcaccccatctttgggaatattgttgacgaagtag
cctatcatgaaaagtatccgactatataccaccttcgcaagaagctggtggactcaaccgataaggcagaccttcggctcatatacctggctctcg
cgcacatgataaagtttcgtggccatttcttgatcgaaggggacctcaacccggataactccgatgtggataaactgttcattcagctcgtccaaa
cctacaatcagctgttcgaggagaaccccatcaatgcatcaggtgtcgacgccaaggcaatactgtctgccagactttcgaagtccagacggctt
gagaatctgatcgctcaattgccaggcgagaagaagaacggcttgttcgggaatctgattgcactgtctctgggcctcacccctaacttcaaaagc
aactttgacctcgccgaggacgcgaagctgcagctgtcaaaggatacatacgatgatgatctggacaatctgctcgcccaaataggtgatcagta
tgccgacctgttcttggctgccaagaatctgtcagacgctatcttgctcagtgacattctgcgggtcaacacggagataaccaaagcgccacttag
cgcctccatgatcaagaggtacgacgagcatcaccaggatctgacccttctgaaggctttggttcgccagcaactccccgagaagtacaaggag
attttctttgaccaatcgaagaatggctacgcagggtacattgatggaggtgcaagtcaggaggaattctacaaattcatcaagcctattctggaa
aagatggacggtacagaggagctgctcgttaaattgaaccgcgaagatttgcttcggaagcagcgtaccttcgacaatggcagcataccgcacc
agatccacctcggtgagctgcatgctatcttgaggaggcaagaggacttctatccgttcctgaaagacaacagagagaagattgaaaagatcct
cacgttccgcattccctactatgtaggtccactcgcacgcgggaactcgcggtttgcgtggatgacacgcaaatccgaggagactatcacgccttg
gaacttcgaagaggtcgtggacaagggtgcgagtgcacagtccttcatcgaaaggatgaccaacttcgataagaatctcccaaatgagaaagtc
ctgcccaagcatagtctcctgtacgaatacttcacggtctacaacgagctgacgaaggtgaaatatgtgacggaggggatgcgcaaaccggcctt
cctgtcaggtgagcagaagaaggccattgtcgatctcttgttcaaaaccaatcggaaggtcactgtgaaacagcttaaagaggactactttaaga
agatcgaatgctttgattctgtggaaatcagcggcgttgaggataggttcaatgcctctcttggcacataccatgacctgttgaaaatcatcaagg
acaaggacttccttgacaacgaggagaacgaggacatcctcgaggacatcgtgctgactctcacgctgtttgaggacagagaaatgatcgagga
gcgccttaagacttatgcgcatctgttcgatgacaaggtcatgaagcagttgaagaggaggagatatacaggttggggaaggctctccaggaag
ctcatcaacggcatccgcgacaagcaatccggcaagactatactggactttctcaaatccgacggttttgcgaatcggaacttcatgcagcttatt
cacgatgactcactgaccttcaaagaagatatccagaaggcccaagtgtcaggtcagggcgatagccttcacgaacacatagccaacctggctg
gatcgccagctataaagaagggcatactgcagacagtgaaggttgtggatgagctggtgaaggtcatgggccgccataagccggagaacatcg
tcatcgagatggcgagggaaaaccagacgactcagaaagggcagaagaactcacgggagcgcatgaagcggatagaggaaggcatcaagga
gcttgggagtcagattctgaaagagcacccagtcgaaaatactcaactccagaacgagaagctgtacctctattacctccagaatgggagagat
atgtacgtcgaccaagagctcgacattaacagactctccgactatgatgtggatcacattgtccctcaatctttcctgaaggacgatagtattgaca
acaaggtccttacgcgctcagacaagaaccgcggaaaatccgacaatgtacccagcgaggaggttgtgaagaagatgaagaactattggaggc
agcttttgaatgctaagctcataacccaacggaaattcgacaatctcacgaaggcagaaaggggcggactgtctgagctcgacaaagccggctt
catcaagcgccagttggttgaaactcgtcagattacgaaacatgtggcccagatactcgattcgcgtatgaatacgaagtatgatgagaatgaca
aacttatcagggaggtaaaggtgatcaccctcaagagcaaactggttagtgacttccggaaggacttccagttttacaaggttcgcgagatcaac
aactaccatcatgcccatgacgcctacctgaacgccgttgttggcactgctctcatcaagaagtatccgaaactggagtctgagtttgtgtacggg
gattacaaggtgtacgacgttaggaagatgatcgcgaagtcagaacaagagatcggcaaggctaccgcgaaatacttcttttactcgaatatcat
gaacttcttcaagacagagatcactctggcgaatggtgaaatccggaagaggcctctgatcgagacaaatggcgaaacaggtgagattgtctgg
gataagggcagggattttgcgactgtgcgtaaggttctcagcatgccccaagtcaacatagtcaagaaaacggaggttcaaaccggtggtttctc

caaggagtccattctccctaagcgcaactccgacaaactgattgcgaggaagaaggattgggatccgaagaaatacggaggctttgatagccct
accgtggcatacagcgtactggtagtggccaaggtggagaagggcaagagcaagaaactgaaaagcgtcaaggaactgcttggaattaccata
atggaaaggtcctcgttcgagaagaatccgatcgacttcctcgaggctaaaggttacaaagaggtgaagaaagacctcattatcaaactgccca
agtattcgcttttcgaattggaaaatggcagaaaacgcatgctggcatctgccggagaactgcagaagggcaacgagctggcattgcccagtaa
gtacgtcaacttcctgtacttggcctcacactatgagaagctgaaggggtcaccagaggacaacgagcagaagcagttgtttgtcgagcagcac
aagcactatcttgatgagatcatagagcagatcagcgaattttccaagcgggtcattcttgcagacgctaacctcgataaggtgctttccgcgtac
aacaagcacagagataagccgataagggaacaagcggaaaacatcatccacctgttcacactgaccaatctgggagccccagcagcctttaag
tacttcgataccactatcgacagaaagcgctacacatcaaccaaggaagtgttggacgctacccttattcaccaatctattacagggctctatgag
acaaggatagatctgtcgcagttgggtggtgac **tctagggctgacccaaagaagaagcgtaaagtc** TGA

*Zea mays* codon-optimized Cas9-NLS with introns (zCas9i) of pAGM51547 and pAGM51559 (introns in yellow)

ATGgacaagaagtacagcattggacttgatattggtacgaactcagttgggtgggccgttatcaccgatgaatacaaggtaccttcgaagaaat
ttaaagtgctgggcaacacagataggcacagcattaagaagaacttgatcggagctctgctctttgactctggagaaaccgcggaggcgacaag
gcttaaacgtactgcgaggagaaggtacactcgcaggaagaacagaatctgttatctccaagagatctttagcaacgagatggcgaag*gtaag*
*gatttttatgatatagtatgcttatgtattttgtactgaaagcatatcctgcttcattgggatattactgaaagcatttaactacatgtaaactc*
*acttgatgatcaataaacttgattttgcag*gttgacgactcgttcttccatcgcctcgaggaatccttcctggtagaggaagataagaaacacga
gcgtcaccccatctttgggaatattgttgacgaagtagcctatcatgaaaagtatccgactatataccaccttcgcaagaagctggtggactcaac
cgataaggcagaccttcggctcatatacctggctctcgcgcacatgataaagtttcgtggccatttcttgatcgaaggggacctcaacccggataa
ctccgatgtggataaactgttcattcagctcgtccaaacctacaatcagctgttcgaggagaaccccatcaatgcatcag*gtaacattccttagtt*
*acctttcttttctttttccatcataagtttatagattgtacatgctttgagatttttctttgcaaacaatctcag*gtgtcgacgccaaggcaatactg
tctgccagactttcgaagtccagacggcttgagaatctgatcgctcaattgccaggcgagaagaagaacggcttgttcgggaatctgattgcactg
tctctgggcctcacccctaacttcaaaagcaactttgacctcgccgaggacgcgaagctgcagctgtcaaaggatacatacgatgatgatctgga
caatctgctcgcccaaatag*gtgctcttgaaattggaactcttcttttgttgtctaaacctatcaatttctttgcggaaatttatttgaagctgtag*
*agttaaaattgagtcttttaaacttttgtag*gtgatcagtatgccgacctgttcttggctgccaagaatctgtcagacgctatcttgctcagtgaca
ttctgcgggtcaacacggagataaccaaagcgccacttagcgcctccatgatcaagaggtacgacgagcatcaccaggatctgacccttctgaa
ggctttggttcgccagcaactccccgagaagtacaaggagattttctttgaccaatcgaagaatggctacgcagggtacattgatggag*gtaagt*
*tgttacttatgattgttttcctctctgctacatgtattttgttgttcatttctgtaagatataagaattgagttttcctctgatgatattattag*gtgc
aagtcaggaggaattctacaaattcatcaagcctattctggaaaagatggacggtacagaggagctgctcgttaaattgaaccgcgaagatttgc
ttcggaagcagcgtaccttcgacaatggcagcataccgcaccagatccacctcggtgagctgcatgctatcttgaggaggcaagaggacttctat
ccgttcctgaaagacaacagagagaagattgaaaagatcctcacgttccgcattccctactatgtag*gttagtatcatatgaagaaatacctag*
*tttcagttgatgaatgctattttctgacctcagttgttctcttttgagaattatttcttttctaatttgcctgatttttctattaattcattag*gtccact
cgcacgcgggaactcgcggtttgcgtggatgacacgcaaatccgaggagactatcacgccttggaacttcgaagaggtcgtggacaagggtgcg
agtgcacagtccttcatcgaaaggatgaccaacttcgataagaatctcccaaatgagaaagtcctgcccaagcatagtctcctgtacgaatacttc
acggtctacaacgagctgacgaaggtgaaatatgtgacggaggggatgcgcaaaccggccttcctgtcag*gtaaatcctggtccacacttttac*
*gataaaaacacaagattttaaactatgaactgatcaataatcattcctaaaagaccacacttttgttttgtttctaaagtaatttttactgttat*
*aacag*gtgagcagaagaaggccattgtcgatctcttgttcaaaaccaatcggaaggtcactgtgaaacagcttaaagaggactactttaagaag
atcgaatgctttgattctgtggaaatcagcggcgttgaggataggttcaatgcctctcttggcacataccatgacctgttgaaaatcatcaaggac
aaggacttccttgacaacgaggagaacgaggacatcctcgaggacatcgtgctgactctcacgctgtttgaggacagagaaatgatcgaggagc
gccttaagacttatgcgcatctgttcgatgacaaggtcatgaagcagttgaagaggaggagatatacag*gtaagaggtcaaaaggtttccgca*
*atgatccctctttttttgtttctctagtttcaagaatttgggtatatgactaacttctgagtgttccttgatgcatatttgtgatgagacaaatgttt*
*gttctatgttttag*gttggggaaggctctccaggaagctcatcaacggcatccgcgacaagcaatccggcaagactatactggactttctcaaatc
cgacggttttgcgaatcggaacttcatgcagcttattcacgatgactcactgaccttcaaagaagatatccagaaggcccaagtgtcaggtcagg
gcgatagccttcacgaacacatagccaacctggctggatcgccagctataaagaagggcatactgcagacagtgaaggttgtggatgagctggt

gaag*gtaagttctgcatttggttatgctccttgcattttaggtgttcgtcgcacttccatttccatgaatagctaagatttttttctctgcattcatt*
*cttcttgcctcagttctaactgtttgtggtatttttgttttaattattgctacag*gtcatgggccgccataagccggagaacatcgtcatcgagatgg
cgagggaaaaccagacgactcagaaagggcagaagaactcacgggagcgcatgaagcggatagaggaaggcatcaaggagcttgggagtca
gattctgaaagagcacccagtcgaaaatactcaactccagaacgagaagctgtacctctattacctccagaatgggagagatatgtacgtcgac
caagagctcgacattaacagactctccgactatgatgtggatcacattgtccctcaatctttcctgaaggacgatagtattgacaacaag*gtaaa*
*gcaactgtgttttaatcaatttcttgtcaggatatatggattataacttaatttttgagaaatctgtagtatttggcgtgaaatgagtttgctttt*
*tggtttctcccgtgttatag*gtccttacgcgctcagacaagaaccgcggaaaatccgacaatgtacccagcgaggaggttgtgaagaagatgaa
gaactattggaggcagcttttgaatgctaagctcataacccaacggaaattcgacaatctcacgaaggcagaaaggggcggactgtctgagctc
gacaaagccggcttcatcaagcgccagttggttgaaactcgtcagattacgaaacatgtggcccagatactcgattcgcgtatgaatacgaagta
tgatgagaatgacaaacttatcagggaggtaaag*gtaaagtttccaactttcctttaccatatcaaactaaagttcgaaactttttatttgatca*
*acttcaaggccacccgatctttctattcctgattaatttgtgatgaatccatattgacttttgatggttacgcag*gtgatcaccctcaagagcaaa
ctggttagtgacttccggaaggacttccagttttacaaggttcgcgagatcaacaactaccatcatgcccatgacgcctacctgaacgccgttgttg
gcactgctctcatcaagaagtatccgaaactggagtctgagtttgtgtacggggattacaaggtgtacgacgttaggaagatgatcgcgaagtca
gaacaagagatcggcaaggctaccgcgaaatacttcttttactcgaatatcatgaacttcttcaagacagagatcactctggcgaatggtgaaat
ccggaagaggcctctgatcgagacaaatggcgaaacag*gtctgtctttcctatttcatatgtttaatcctaggaatttgatcaattgattgtatgt*
*atgtcgatcccaagactttcttgttcacttatatcttaactctctctttgctgtttcttgcag*gtgagattgtctgggataagggcagggattttgcg
actgtgcgtaaggttctcagcatgccccaagtcaacatagtcaagaaaacggaggttcaaaccggtggtttctccaaggagtccattctccctaag
cgcaactccgacaaactgattgcgaggaagaaggattgggatccgaagaaatacggaggctttgatagccctaccgtggcatacagcgtactgg
tagtggccaaggtggagaagggcaagagcaagaaactgaaaagcgtcaaggaactgcttggaattaccataatggaaaggtcctcgttcgaga
agaatccgatcgacttcctcgaggctaaaggtaaaatattggatgccagacgatattctttcttttgatttgtaactttttcctgtcaaggtcgataa
attttatttttttggtaaaaggtcgataattttttttggagccattatgtaattttcctaattaactgaaccaaaattatacaaaccaggttacaaag
aggtgaagaaagacctcattatcaaactgcccaagtattcgcttttcgaattggaaaatggcagaaaacgcatgctggcatctgccggagaactg
cagaagggcaacgagctggcattgcccagtaagtacgtcaacttcctgtacttggcctcacactatgagaagctgaaggggtcaccagaggaca
acgagcagaagcagttgtttgtcgagcagcacaagcactatcttgatgagatcatagagcagatcagcgaattttccaagcgggtcattcttgca
gacgctaacctcgataag*gtaaggacttctcatgaatattagtggcagattagtgttgttaaagtctttggttagataatcgatgcctcctaat*
*tgtccatgttttactggttttctacaattaaag*gtgctttccgcgtacaacaagcacagagataagccgataagggaacaagcggaaaacatca
tccacctgttcacactgaccaatctgggagccccagcagcctttaagtacttcgataccactatcgacagaaagcgctacacatcaaccaaggaa
gtgttggacgctacccttattcaccaatctattacagggctctatgagacaaggatagatctgtcgcagttgggtggtgac
**tctagggctgacccaaagaagaagcgtaaagtc** TGA

*Zea mays* codon-optimized Flag tag-NLS-Cas9-NLS with introns (zCasio) of pAGM51561 (Flag tag-NLS in green, introns in yellow, intron mutations in red underlined, and N-terminal NLS in red)

ATGgactataaggaccacgacggagactacaaggatcatgatattgattacaaagacgatgacgataagatggccccaaagaagaagcgga
aggtcggtatccacggagtcccagcagccgacaagaagtacagcattggacttgatattggtacgaactcagttgggtgggccgttatcaccgat
gaatacaaggtaccttcgaagaaatttaaagtgctgggcaacacagataggcacagcattaagaagaacttgatcggagctctgctctttgactc
tggagaaaccgcggaggcgacaaggcttaaacgtactgcgaggagaaggtacactcgcaggaagaacagaatctgttatctccaagagatctt
tagcaacgagatggcgaag*gtaaggattttatgatatactatgcttatgtattttgtactgaaagcatatcctgcttcattgggatattactga*
*aagcatttaactacatgtaaactcacttgatgatcaataaacttgattttgcag*gttgacgactcgttcttccatcgcctcgaggaatccttcctg
gtagaggaagataagaaacacgagcgtcaccccatctttgggaatattgttgacgaagtagcctatcatgaaaagtatccgactatataccacct
tcgcaagaagctggtggactcaaccgataaggcagaccttcggctcatatacctggctctcgcgcacatgataaagtttcgtggccatttcttgatc
gaaggggacctcaacccggataactccgatgtggataaactgttcattcagctcgtccaaacctacaatcagctgttcgaggagaaccccatcaa
tgcatcag*gtaacattccttagttacctttcttttctttttccatcataagtttatagattgtacatgctttgagatttttctttgcaaacaatctcag*
gtgtcgacgccaaggcaatactgtctgccagactttcgaagtccagacggcttgagaatctgatcgctcaattgccaggcgagaagaagaacgg
cttgttcgggaatctgattgcactgtctctgggcctcacccctaacttcaaaagcaactttgacctcgccgaggacgcgaagctgcagctgtcaaa

ggatacatacgatgatgatctggacaatctgctcgcccaaatag*gt**aa**tcttgaaattggaactcttcttttgttgtctaaacctatcaatttcttt*
*gcggaaatttatttgaagctgtagagttaaaattgagtcttttaaacttttgtag*gtgatcagtatgccgacctgttcttggctgccaagaatctg
tcagacgctatcttgctcagtgacattctgcgggtcaacacggagataaccaaagcgccacttagcgcctccatgatcaagaggtacgacgagca
tcaccaggatctgacccttctgaaggctttggttcgccagcaactccccgagaagtacaaggagattttctttgaccaatcgaagaatggctacgc
agggtacattgatggag*gtaagttgttacttatgattgttttcctctctgctacatgtattttgttgttcatttctgtaagatataagaattgagttt*
*tcctctgatgatattattag*gtgcaagtcaggaggaattctacaaattcatcaagcctattctggaaaagatggacggtacagaggagctgctcg
ttaaattgaaccgcgaagatttgcttcggaagcagcgtaccttcgacaatggcagcataccgcaccagatccacctcggtgagctgcatgctatct
tgaggaggcaagaggacttctatccgttcctgaaagacaacagagagaagattgaaaagatcctcacgttccgcattccctactatgtag*gttag*
*tatcatatgaagaaatacctagtttcagttgatgaatgctattttctgacctcagttgttctcttttgagaattatttcttttctaatttgcctgattt*
*ttctattaattcattag*gtccactcgcacgcgggaactcgcggtttgcgtggatgacacgcaaatccgaggagactatcacgccttggaacttcga
agaggtcgtggacaagggtgcgagtgcacagtccttcatcgaaaggatgaccaacttcgataagaatctcccaaatgagaaagtcctgcccaag
catagtctcctgtacgaatacttcacggtctacaacgagctgacgaaggtgaaatatgtgacggaggggatgcgcaaaccggccttcctgtcag*g*
*taaatcctggtccacacttttacgataaaaacacaagattttaaactatgaactgatcaataatcattcctaaaagaccacacttttgttttgt*
*ttctaaagtaatttttactgttataacag*gtgagcagaagaaggccattgtcgatctcttgttcaaaaccaatcggaaggtcactgtgaaacagc
ttaaagaggactactttaagaagatcgaatgctttgattctgtggaaatcagcggcgttgaggataggttcaatgcctctcttggcacataccatg
acctgttgaaaatcatcaaggacaaggacttccttgacaacgaggagaacgaggacatcctcgaggacatcgtgctgactctcacgctgtttgag
gacagagaaatgatcgaggagcgccttaagacttatgcgcatctgttcgatgacaaggtcatgaagcagttgaagaggaggagatatacag*gt*
*aagaggtcaaaaggtttccgcaatgatccctctttttttgtttctctagtttcaagaatttgggtatatgactaacttctgagtgttccttgatgc*
*atatttgtgatgagacaaatgtttgttctatgttttag*gttggggaaggctctccaggaagctcatcaacggcatccgcgacaagcaatccggca
agactatactggactttctcaaatccgacggttttgcgaatcggaacttcatgcagcttattcacgatgactcactgaccttcaaagaagatatcca
gaaggcccaagtgtcaggtcagggcgatagccttcacgaacacatagccaacctggctggatcgccagctataaagaagggcatactgcagac
agtgaaggttgtggatgagctggtgaag*gtaagttctgcatttggttatgctccttgcattttaggtgttcgtcgcacttccatttccatgaatagc*
*taagattttttttctctgcattcattcttcttgcctcagttctaactgtttgtggtatttttgttttaattattgctacag*gtcatgggccgccataagc
cggagaacatcgtcatcgagatggcgagggaaaaccagacgactcagaaagggcagaagaactcacgggagcgcatgaagcggatagagga
aggcatcaaggagcttgggagtcagattctgaaagagcacccagtcgaaaatactcaactccagaacgagaagctgtacctctattacctccag
aatgggagagatatgtacgtcgaccaagagctcgacattaacagactctccgactatgatgtggatcacattgtccctcaatctttcctgaaggac
gatagtattgacaacaag*gtaaagcaactgtgttttaatcaatttcttgtcaggatatatggattataacttaatttttgagaaatctgtagtat*
*ttggcgtgaaatgagtttgctttttggtttctcccgtgttatag*gtccttacgcgctcagacaagaaccgcggaaaatccgacaatgtacccagc
gaggaggttgtgaagaagatgaagaactattggaggcagcttttgaatgctaagctcataacccaacggaaattcgacaatctcacgaaggcag
aaaggggcggactgtctgagctcgacaaagccggcttcatcaagcgccagttggttgaaactcgtcagattacgaaacatgtggcccagatactc
gattcgcgtatgaatacgaagtatgatgagaatgacaaacttatcagggaggtaaag*gtaaagtttccaactttcctttaccatatcaaactaa*
*agttcgaaactttttatttgatcaacttcaaggccacccgatctttctattcctgattaatttgtgatgaatccatattgacttttgatggttacgc*
*ag*gtgatcaccctcaagagcaaactggttagtgacttccggaaggacttccagttttacaaggttcgcgagatcaacaactaccatcatgcccat
gacgcctacctgaacgccgttgttggcactgctctcatcaagaagtatccgaaactggagtctgagtttgtgtacggggattacaaggtgtacgac
gttaggaagatgatcgcgaagtcagaacaagagatcggcaaggctaccgcgaaatacttcttttactcgaatatcatgaacttcttcaagacaga
gatcactctggcgaatggtgaaatccggaagaggcctctgatcgagacaaatggcgaaacag*gtctgtctttcctatttcatatgtttaatcctag*
*gaatttgatcaattgattgtatgtatgtcgatcccaagactttcttgttcacttatatcttaactctctctttgctgtttcttgcag*gtgagattgtct
gggataagggcagggattttgcgactgtgcgtaaggttctcagcatgccccaagtcaacatagtcaagaaaacggaggttcaaaccggtggtttc
tccaaggagtccattctccctaagcgcaactccgacaaactgattgcgaggaagaaggattgggatccgaagaaatacggaggctttgatagcc
ctaccgtggcatacagcgtactggtagtggccaaggtggagaagggcaagagcaagaaactgaaaagcgtcaaggaactgcttggaattacca
taatggaaaggtcctcgttcgagaagaatccgatcgacttcctcgaggctaaaggtaaaatattggatgccagacgatattctttcttttgatttgt
aactttttcctgtcaaggtcgataaattttattttttttggtaaaaggtcgataattttttttggagccattatgtaattttcctaattaactgaaccaa
aattatac**tttg**caggttacaaagaggtgaagaaagacctcattatcaaactgcccaagtattcgcttttcgaattggaaaatggcagaaaacgc
atgctggcatctgccggagaactgcagaagggcaacgagctggcattgcccagtaagtacgtcaacttcctgtacttggcctcacactatgagaa
gctgaaggggtcaccagaggacaacgagcagaagcagttgtttgtcgagcagcacaagcactatcttgatgagatcatagagcagatcagcga
attttccaagcgggtcattcttgcagacgctaacctcgataag*gtaaggacttctcatgaatattagtggcagattagtgttgttaaagtctttg*

*gttagataatcgatgcctcctaattgtccatgttttactggttttctacaatta**c**ag*gtgctttccgcgtacaacaagcacagagataagccgata agggaacaagcggaaaacatcatccacctgttcacactgaccaatctgggagccccagcagcctttaagtacttcgataccactatcgacagaa agcgctacacatcaaccaaggaagtgttggacgctacccttattcaccaatctattacagggctctatgagacaaggatagatctgtcgcagttgg gtggtgac **tctagggctgacccaaagaagaagcgtaaagtc** TGA

**Figure S13: Cloning of a Guide RNA in a MoClo level 1 construct.** (A) Primer design. A PCR product containing the guide RNA sequences is amplified using plasmid pAGM9037 as a template using primers critarx and critarev. Primer critarx contains a BsaI site (ggtctc) followed by 4 nucleotides overlapping with the end of the Arabidopsis U6 promoter (attg), the variable sequence of the target site (20 n in blue), and by 19 nucleotides of the constant region of the guide RNA. Primer critarev is a primer binding a sequence in the vector backbone downstream of the terminator of the guide RNA. (B) Cloning of the PCR product in a level 1 MoClo vector. The amplified product is cloned together with a promoter of choice (pICSL01009 or pAGM38869) in a MoClo level 1 vector (for example pICH47751 for position 3) using BsaI and ligase. The amplified part of the construct is sequenced with primer l1f (in flanking vector sequences).

A

target site starts with a g

(PAM)

g nnn nnn nnn nnn nnn nnn n ngg

critarx     tt ggtctc a attg nnn nnn nnn nnn nnn nnn n gttttagagctagaaatag

target site starts does not with a g

n nnn nnn nnn nnn nnn nnn n ngg

critarx     t ggtctc a attg n nnn nnn nnn nnn nnn nnn n gttttagagctagaaatag

critarev     atgtacggccagcaacgtcg     (reverse primer, always the same)

B

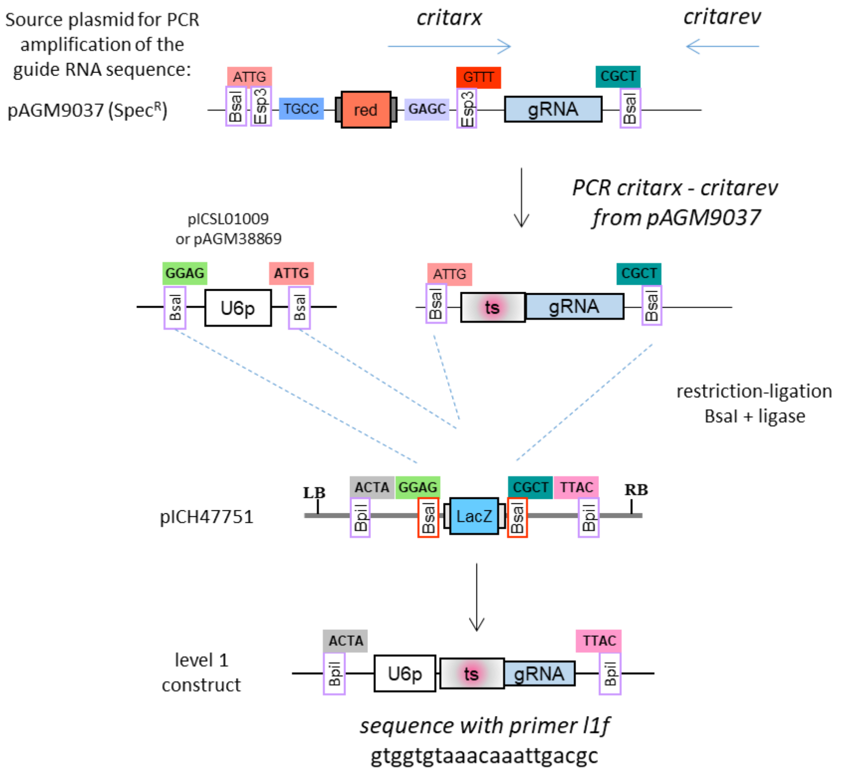

**Figure S14: Construction of Cas9 constructs containing 1 to 4 guide RNAs.** The cloning strategy is done in two steps. First, the chosen number of guide RNAs (one to four) are cloned in level 1 vectors position 3 to 6 in vectors pICH47751, pICH47761, pICH47772 and pICH47781, respectively, as described in Figure S13. The final level M vector is made by subcloning fragments from pAGM35131 (transformation marker), pAGM52323 (Cas9 expression cassette), the chosen number of guide RNAs from level 1 constructs, and the appropriate level M end-linker, in level M vector pAGM8031 using BpiI and ligase.

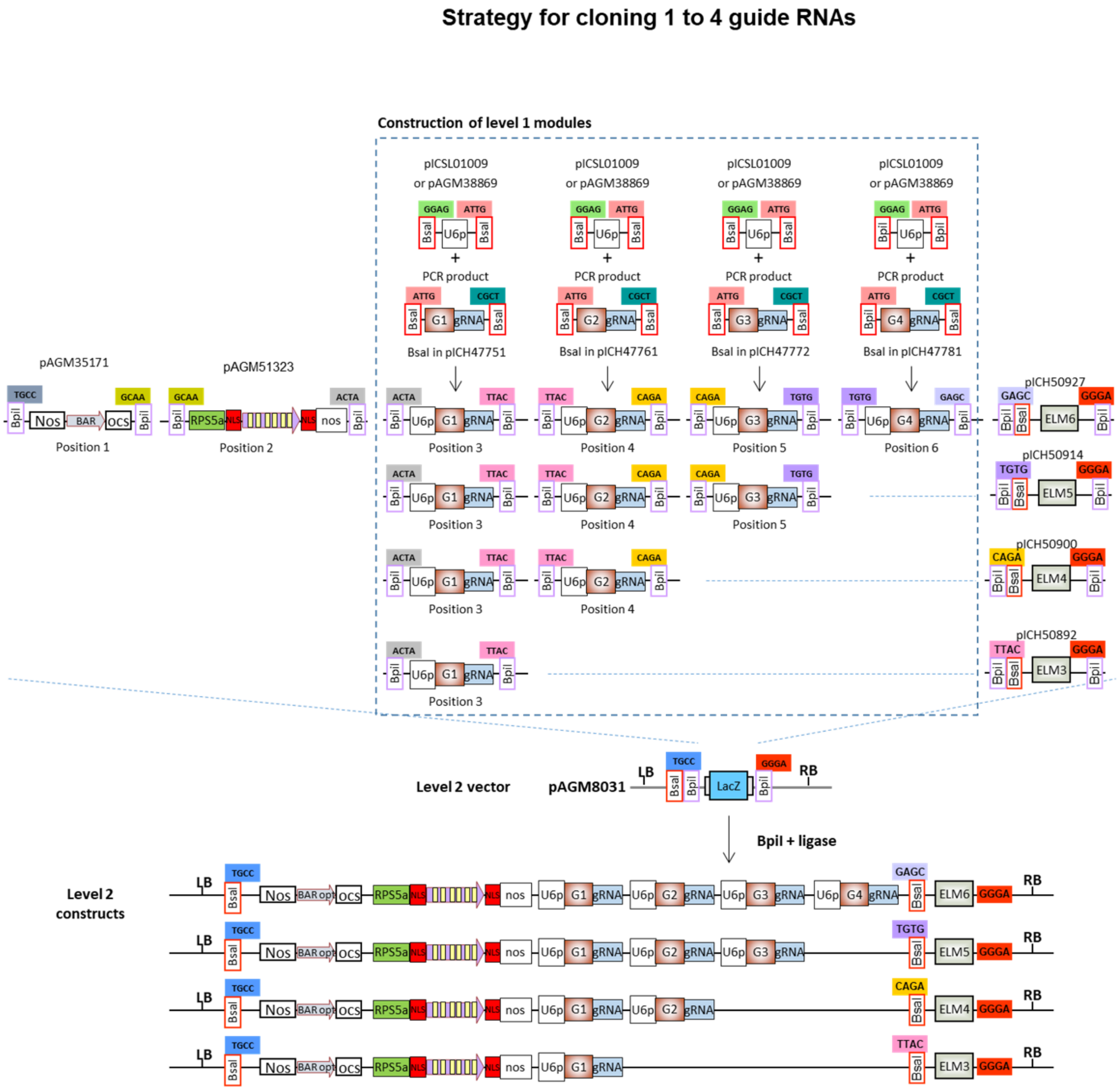

**Figure S15a: Construction of Cas9 constructs containing 5 to 10 guide RNAs.** As in the previous example, the first step consists of cloning guide RNAs in level 1 constructs. In this case, they are cloned in level 1 vectors for position 7, 1, 2, 3, 4, and 5 as described in Figure S13 using BsaI and ligase. Then the selected number of level 1 guide RNA constructs are subcloned in level M vector pAGM8093 with the appropriate end-linker using BpiI and ligase.

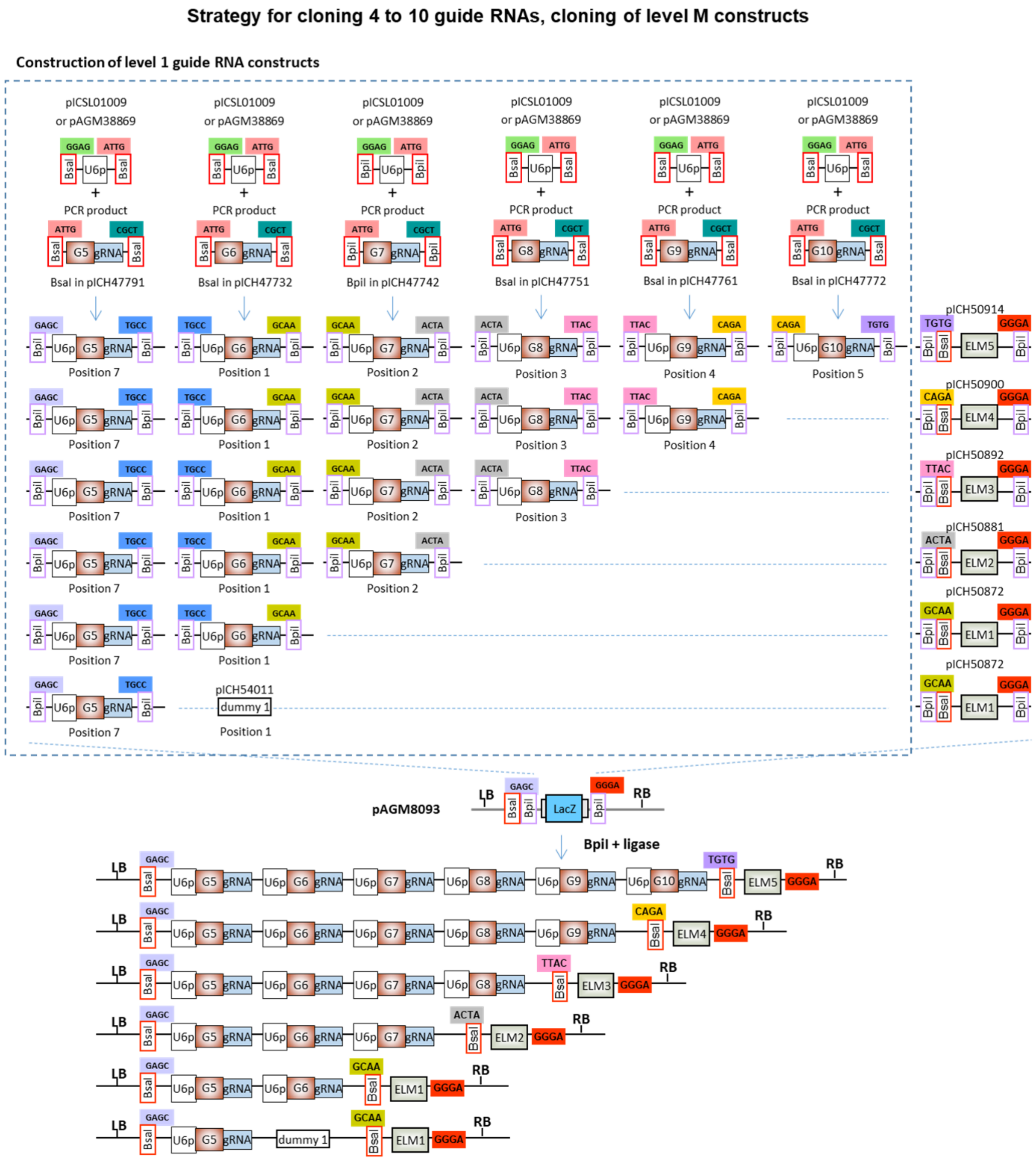

**Figure S15b: Construction of Cas9 constructs containing 5 to 10 guide RNAs.** The final step is the assembly of two level M constructs, the first one containing 4 guide RNAs (made in Figure S14), and the second one containing 1 to 6 guide RNAs (made in Figure S15a) with the appropriate end-linker in pICH75322 using BsaI and ligase.

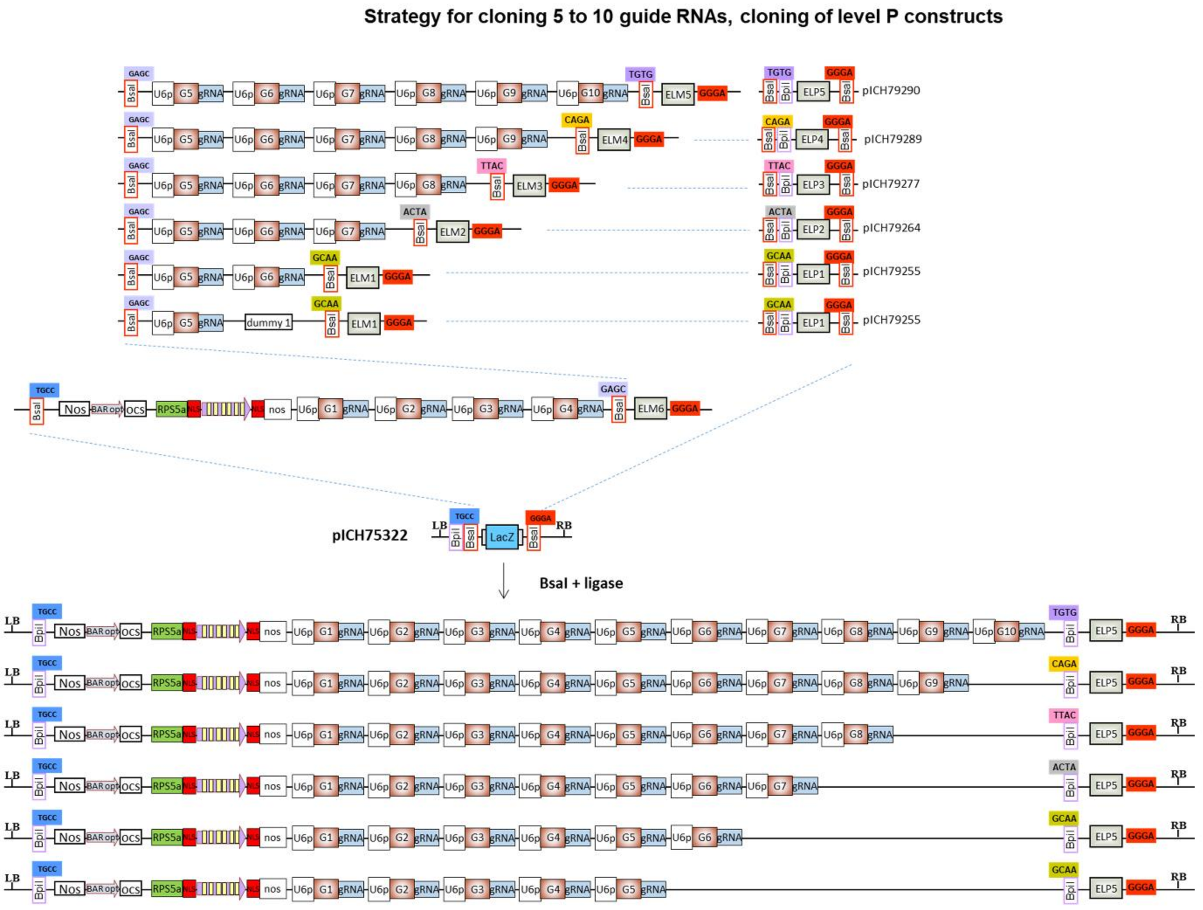

**Figure S16: Cloning 1 to 6 guide RNAs in level 2 vectors already containing Cas9.** As a first step guide RNAs are cloned in level1 vectors and sequenced. Level 2 constructs are made using the selected level 1 constructs, the corresponding end-linker, and a selected level 2 cloning vector.

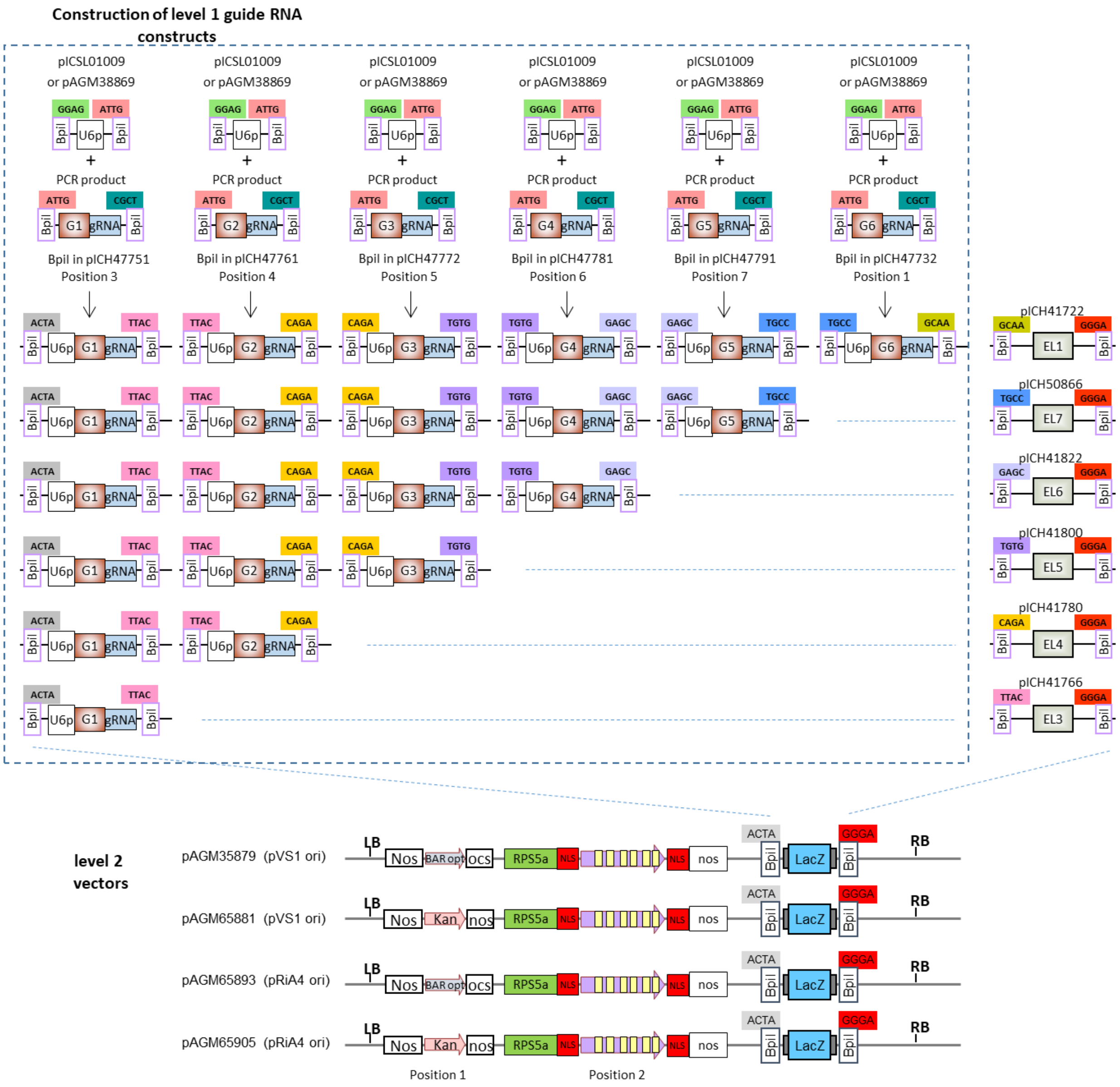
